# Condensate Material Properties Influence Cargo Selection for Neuronal Extrusion via Large Extracellular Exopher Vesicles

**DOI:** 10.64898/2026.08.17.745362

**Authors:** Mayur Barai, Edward Chuang, Shuo-Lin Weng, Jeetain Mittal, Monica Driscoll, Benjamin S. Schuster

**Author notes:** Co-first authors contributed equally to this work (Mayur Barai, Edward Chuang).

## Abstract

Neurons can maintain proteostasis by extruding protein aggregates and damaged organelles via large extracellular vesicles called exophers. However, the biophysical rules governing exopher cargo selection remain poorly understood. Here, we discovered that biomolecular condensates can localize to exophers and investigated whether condensate material properties—ranging from liquids to dynamically arrested gels—determine their recruitment to exophers. By reprogramming interactions in a multidomain protein, we tuned condensate properties and linked them to neuronal exopher recruitment using a multidisciplinary approach combining *in vivo* dynamics, *in vitro* material properties, and molecular dynamics simulations. Our findings reveal that gel-like condensates with slower dynamics and higher viscoelasticity localize to exophers more avidly than dynamic liquid-like condensates, demonstrating a strong correlation between condensate dynamics and exopher recruitment. This work lays the framework for investigating the biophysical determinants of exopher cargo selection, providing critical insights into the influence of proteome material state on neuronal protein quality control.

**Graphical Abstract:** 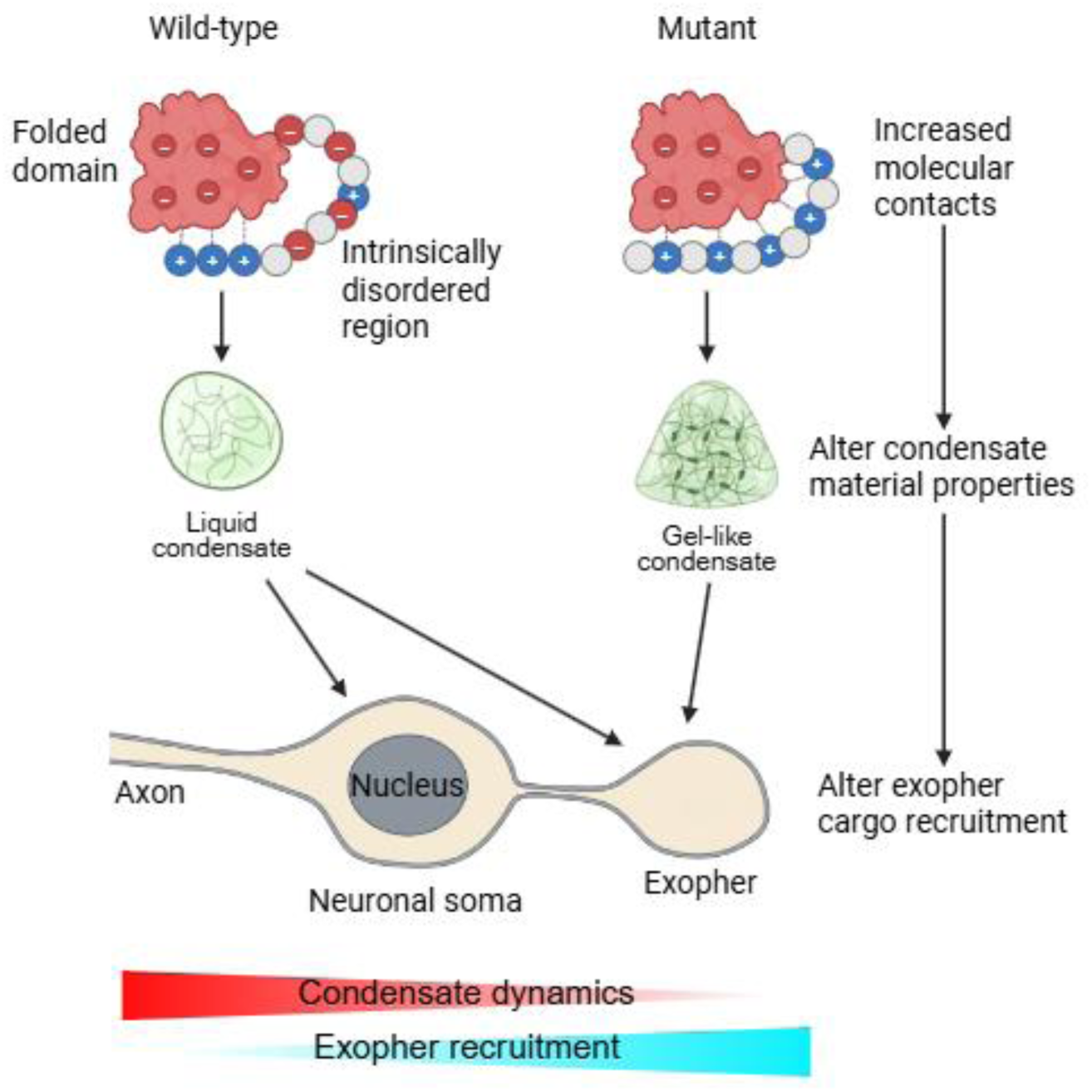

## Introduction

Proteostasis anchors cellular function and survival. To preserve a healthy proteome, cells utilize a network of quality control mechanisms, including the use of molecular chaperones for protein folding, the ubiquitin-proteasome system for targeted degradation, and autophagy for bulk clearance of damaged or misfolded proteins^1,2^. Neurons, as long-lived post-mitotic cells, are particularly sensitive to proteostasis compromise, a vulnerability underscored by the pathological accumulation of misfolded protein aggregates that are typically found in degenerating neurons in neurodegenerative diseases^3,4^. Defining mechanistic details of the multi-faceted cellular systems for protein quality control is thus a central focus in the quest to maintain nervous system health in aging and disease.

Recent discoveries have revealed a previously unrecognized mechanism by which cells, particularly neurons, manage proteostatic stress: the production of large extracellular vesicles, called exophers, to eliminate cellular components. Exophers, which are comparable in size to neuronal soma, facilitate the removal of protein aggregates and damaged organelles for remote degradation by neighboring cells^5,6,7^. In *C. elegans,* the genesis of exophers can be directly visualized in the context of a living adult nervous system (**Fig. 1A**). Prior work in *C. elegans* demonstrated preferential recruitment of overexpressed mCherry and mutant Huntingtin aggregates (Htt-polyQ128) as cargo^5,8^, and established that neuronal exopher biogenesis is enhanced under proteostress^5,9^.

**Figure 1:**
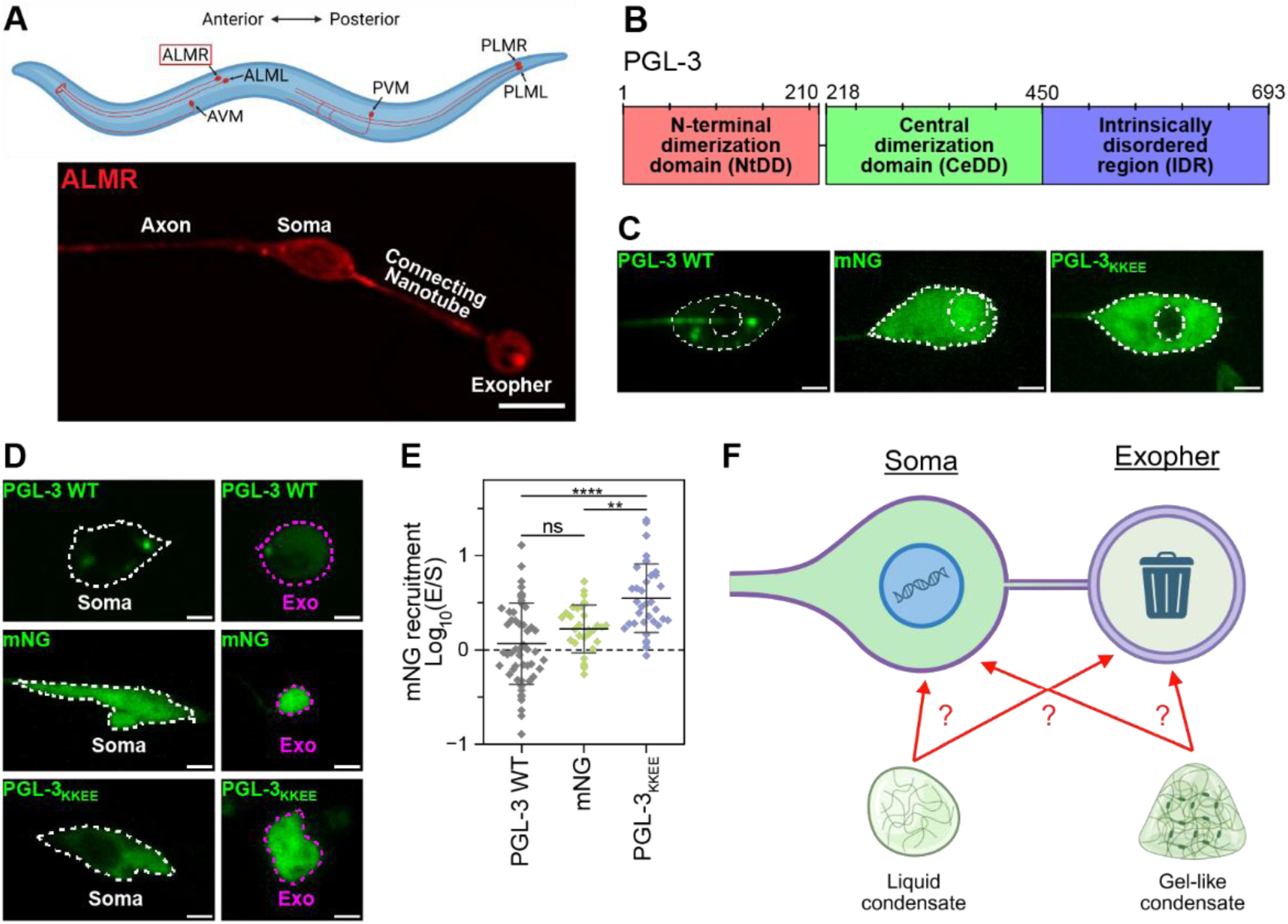
PGL-3 forms foci in TRNs and gets extruded in exophers. **(A)** Cartoon illustrating the *C. elegans* TRN body positions. The six mechanosensory TRNs are ALML, ALMR AVM, PVM, PLML, and PLMR (<u>A</u>nterior/<u>P</u>osterior, <u>L</u>ateral/<u>V</u>entral, <u>M</u>icrotubule, <u>L</u>eft/<u>R</u>ight). Below, an image showing an ALMR exopher in strain bzIs166[*Pmec-4::mCherry*], which ectopically expresses mCherry in the TRNs. mCherry labels the axon, soma, exopher, and thin connecting nanotube between them. Scale bar: 5 μm. mCherry brightness masked and adjusted to highlight all features. The connecting nanotube is significantly dimmer than the soma, which is dimmer than the exopher. **(B)** Domain architecture map for PGL-3. **(C)** Representative ALM neurons from strains expressing PGL-3 WT (ZB5916), mNG (ZB5959), and PGL-3_KKEE_ (ZB5960) in day 2 adults (**Table S2**). PGL-3 WT forms distinct foci while PGL-3_KKEE_ and mNG are diffuse. Soma and nucleus are outlined in white. Scale bar: 2 μm. **(D)** During exopher event, PGL-3 WT foci and diffuse mNG and PGL-3_KKEE_ protein are found in soma and exopher compartments. Day 2 adults fasted for 3 hours, scale bar: 2 μm, soma outlined in white, exopher outlined in magenta. **(E)** PGL-3_KKEE_ localize to exophers more than PGL-3 WT and mNG. Log ratio of mean fluorescence intensity in exopher versus soma of images represented in (D). Zero values represent equal distribution between exopher and soma, positive values indicate more signal in exopher, and negative values indicate more signal in soma. Data represents mean ± S.D. from n=32-51 images per strain. One-way ANOVA and Tukey’s MCT (**p<0.01, ****p<0.0001). **(F)** Cartoon illustrating the proposed question addressed in this study. Are condensates recruited to exophers as cargo? And if so, do the material properties of the condensates determine their recruitment as cargo?

Exopher-like vesicle formation has been reported in multiple *C. elegans* neuronal types^5,10^ and in muscle^11^. In mammalian systems, exopher-like vesicles can be produced by astrocytes and glia^12,13^, axons of the mouse optic nerve head^14,15^, mouse cardiomyocytes^16^, and in human kidney epithelia^17^ or podocytes^18^, suggesting that large extracellular vesicle production is a conserved stress response that can be activated in a range of cell types.

How the material state of the proteome influences neuronal protein quality control is poorly understood. Proteins can undergo phase separation, a process that drives the formation of membraneless compartments called biomolecular condensates, which organize and regulate various cellular processes^19^. Neuronal proteins undergo phase separation to form diverse compartments including neuronal synaptic vesicle clusters^20,21^, RNA-containing granules^22,23^, active zones^24,25^, and postsynaptic densities^26,27^. Functional biomolecular condensates can span a continuum of material properties from dynamic, liquid-like droplets to dynamically arrested, gel-like assemblies^28–32^. Dysregulation of phase separation has been proposed to drive protein aggregation and the development of neurodegenerative diseases. Critically, mutations in aggregation-prone proteins that drive early onset of neurodegenerative diseases can accelerate the liquid-to-solid transition of condensates^32–34^. Solidified condensates can take different forms: while gel-like states exhibit slow dynamics and limited internal protein diffusion relative to liquid condensates^35–38^, aggregation often indicates denaturation and formation of irreversible fibrils.

*C. elegans* exophers can recruit protein aggregates, but whether exophers can recruit biomolecular condensates is unknown. This question is significant, as the extrusion of exopher cargo for remote degradation can promote the health of the proteostressed neuron^5,7^, but conversely, the persistence of deleterious cargo in the sending or receiving cells may underlie the systemic spread of pathology in neurodegenerative disease. We hypothesized that 1) protein condensates could be recruited as exopher cargo, and 2) condensate selection into exophers might be sensitive to condensate material properties. Specifically, we hypothesized that the reduced dynamics of gel-like condensates—as compared to highly dynamic liquid droplets— would favor recruitment to exophers, paralleling the known recruitment of solid aggregates.

Linking condensate material properties to biological functions remains a frontier in the field, as quantitatively measuring condensate properties within living animals is a significant challenge. Recent publications have integrated *in vitro* characterization with qualitative and quantitative *in vivo* measurements to bridge the gap between reconstituted and cellular environments^39–41^. Here we take a similar approach. We utilize PGL-3 as a model protein because of its known ability to form liquid-like condensates *in vitro* and in native *C. elegans* germline^42–46^. PGL-3 also self-assembles into foci when ectopically expressed in somatic cells^47^. By ectopically expressing PGL-3 in *C. elegans* touch receptor neurons (TRNs), we created a tractable system to link mesoscopic material properties to neuronal exopher cargo selection.

We employed a protein engineering approach to rationally tune interactions between the folded domains and the intrinsically disordered region (IDR) of PGL-3 (**Fig. 1B**) to modulate the resulting condensate material properties. We leveraged two complementary strategies to design PGL-3 variants: native IDR mutagenesis and swapping the IDR with artificial intrinsically disordered polypeptide (AIDP) sequences^48^. We further utilized molecular dynamics simulations to identify interactions between PGL-3 domains and to provide mechanistic insights into our experimental observations.

Using these approaches, we discovered that the material properties of condensates can impact their recruitment or localization into exophers—condensates with more gel-like properties are recruited more readily into exophers. We propose that exophers can preferentially recruit condensates on the basis of material properties, as one of the potential factors in exopher cargo sorting. Our work also demonstrates a strategy for engineering biomolecular condensates that modulate their formation and biophysical properties *in vitro* and *in vivo*, aimed at advancing our understanding of cellular functions involving condensates.

## Results

### Neuronally expressed PGL-3 forms foci in soma and in exophers

The *C. elegans* TRN exopher model enables *in vivo* visualization of fluorescent cargo, with exophers expelled from TRNs easily identified as soma-sized fluorescent entities proximal to the sending neuron^5^ (**Fig. 1A**). To develop a system with a model cargo protein that can undergo phase separation and can potentially be recruited into exophers in TRNs, we chose to engineer the PGL-3 protein.

We first asked whether PGL-3 forms discrete foci when ectopically expressed in TRNs. To visualize PGL-3 *in vivo*, we tagged the PGL-3 N-terminus with mNeonGreen (mNG), a monomeric green fluorescent protein. We used the TRN-specific promoter *mec-7* to drive high expression in TRNs from a single-copy insertion of *mNG::pgl-3*. To assist our exopher studies, we included the TRN mCherry allele *bzIs166[Pmec-4::mCherry]* from strain ZB4065, which modestly induces exopher production and enables us to identify neuronal soma and exophers based on mCherry fluorescence (**Fig. 1A**)^5^. For simplicity, hereafter we will refer to the protein mNG::PGL-3 and the strain expressing neuronal mNG::PGL-3 and mCherry (strain ZB5916) as PGL-3 WT (**Table S2**).

We found that ectopic expression of PGL-3 WT is well-tolerated, with no evidence of developmental or fertility compromise in transgenic animals. As the anterior lateral microtubule (ALM) neurons produce exophers at relatively high rates^5^, we examined ALM neurons of day 2 adults and found that PGL-3 WT forms small foci with an average number of ∼2.5 foci per soma and an average size of ∼0.3 μm^2^ (**Fig. 1C, SI Fig. 1**).

PGL-3 features two dimerization domains—an N-terminal dimerization domain (NtDD) and a central dimerization domain (CeDD) (**Fig. 1B**)^45,49^. Previous studies have shown that PGL-3 forms dimers via the two dimerization domains, resulting in phase separation, and that mutations (K126E, K129E; referred to as KKEE) at the dimerization interface of NtDD abrogate phase separation^49,50^. To test whether PGL-3 WT foci formation in TRN is also dependent on PGL-3 dimerization, we expressed mNG without PGL-3 (strain ZB5959) and mNG-tagged PGL-3_KKEE_ (strain ZB5960); both strains used the *mec-7* promoter in the background of the TRN mCherry allele (**Table S2**). We found that both mNG and PGL-3_KKEE_ remain diffuse in the soma (**Fig. 1C**), indicating that PGL-3 foci formation in TRNs likely reflects the intrinsic phase separation capacity of the PGL-3 sequence and that dimerization is essential for PGL-3 assembly into foci, consistent with the literature^49^. Our observations also support that *in vivo* foci formation is not driven by the mNG tag.

Having verified that ectopically expressed PGL-3 forms foci, we next tested whether PGL-3 is localized into exophers. We found that PGL-3 WT foci are present in both the soma and the exophers produced by ALM neurons in day 2 fasted adults (**Fig. 1D**; transient food withdrawal is a stimulus that increases exophergenesis^9^), confirming that PGL-3 can be exopher cargo.

We quantified the relative localization of the three proteins by comparing the log of the ratio of mNG fluorescence in the exopher vs. soma (Log_10_(E/S)) (**Fig. 1E**). We found that PGL-3 WT is evenly distributed between the exopher and soma (Log_10_(E/S) = 0.067±0.43). Interestingly, PGL-3_KKEE_ is significantly enriched and mNG slightly enriched in exophers (possibly mNG more than GFP^5^). Because both PGL-3_KKEE_ and mNG are not expected to form biomolecular condensates themselves, we note that multiple factors likely contribute to exopher cargo selection in the native context. Since our aim here was to understand biomolecular condensate selection into exophers, we chose to build upon our finding that PGL-3 WT can form foci that distribute to soma and exophers in TRNs, seeking to address whether liquid or gel-like condensates are differentially localized to exophers (**Fig. 1F**).

### PGL-3 WT foci have liquid-like dynamics in the *C. elegans* TRNs

Having established that PGL-3 WT forms foci that are detectable in both soma and exophers, we addressed whether the PGL-3 WT foci in the TRNs are indeed liquid condensates, similar to the PGL-3 liquid component of P granules in embryos and PGL-3 condensates *in vitro*^42^. We used two complementary techniques to assess PGL-3 properties *in vivo*: Fluorescence Recovery After Photobleaching (FRAP) and Fluorescence Correlation Spectroscopy (FCS). Instead, we conducted FRAP by photobleaching the entire focus in the soma due to small size of the foci formed, and measured the fluorescence recovery within the bleached region as protein molecules exchange between the focus and cytoplasm. FRAP data from 20 anterior TRNs (ALMR, ALML, or AVM) in day 2 adults indicate that 60% of fluorescence intensity in the foci is recovered within 10 s after bleaching (**Fig. 2A**). Our data indicate that a) PGL-3 WT protein dynamically exchanges between foci and cytoplasm, and b) the foci are not membrane-bound.

**Figure 2:**
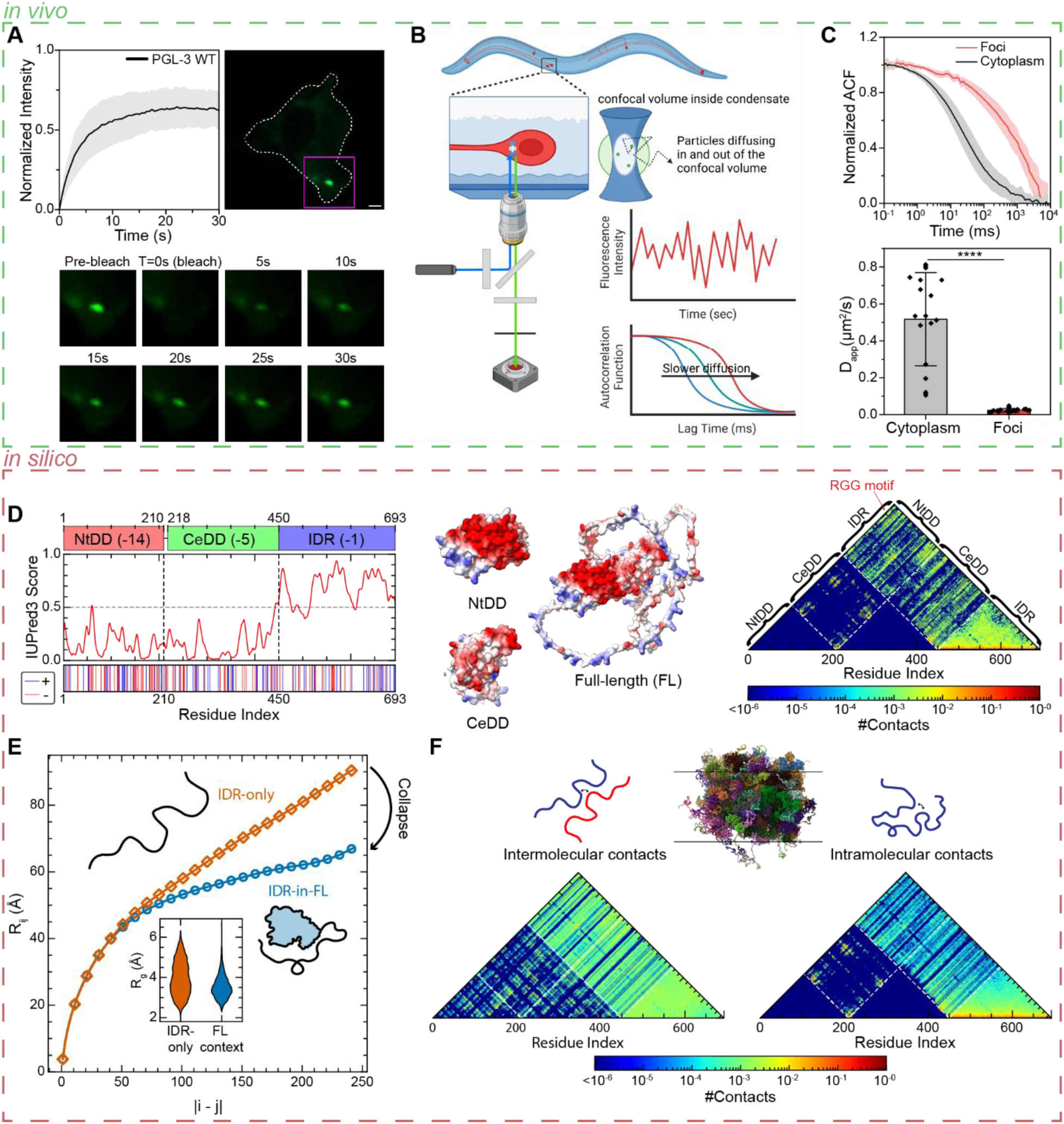
PGL-3 forms dynamic condensates in TRNs and can be modeled by molecular dynamics simulations. **(A) (Top left)** Normalized FRAP recovery traces of PGL-3 WT foci formed in anterior TRN soma show dynamic nature. Data represents mean ± S.D. from n=19 foci from distinct neurons. **(Top right)** Representative image of PGL-3 WT focus in ALMR soma (white dashed line). Scale bar: 1 μm. Box denotes inset for the time lapse images below. **(B)** Schematic illustrating FCS measurements in *C. elegans* TRNs. Confocal laser targets TRN soma just below cuticle. As fluorescent proteins diffuse in and out of the confocal volume, the microscope measures fluctuations in fluorescence intensity. Confocal volume is within cytoplasm or PGL-3 WT focus to measure diffusion of fluorescent protein in each compartment. Autocorrelation of fluorescence intensity data yields autocorrelation function (ACF). Normalized ACF decay rate is inversely proportional to diffusion time of fluorescent proteins. **(C) (Top)** Normalized ACF of PGL-3 WT in cytoplasm and foci in anterior TRN soma showing differences in macromolecular diffusion. **(Bottom)** ACF data are fitted to anomalous diffusion model to estimate apparent diffusion coefficients (D_app_) between cytoplasm and foci. Data represents mean ± SD (foci: n=25, cytoplasm n=18). Welch’s t-test (****p<0.0001). **(D)** Sequence features and charge complementarity of PGL-3. **(Left)** IUPred3 disorder prediction score (line) and charge barcode (blue(+); red(-)). Domains NtDD (residues 1–210, net charge −14), CeDD (residues 218–450, net charge −5), and C-terminal IDR (residues 451–693, net charge −1) are defined using AlphaFold3-predicted structure and IUPred3 Score. **(Middle)** Electrostatic surface potential maps (blue(+); red(-)) of NtDD and CeDD, showing strongly acidic surfaces of both folded domains are poised to interact electrostatically with the basic IDR and highly basic C-terminal tail. **(Right)** Intramolecular contact map derived from PGL-3 WT single-chain coarse-grained (CG) simulation, revealing electrostatic interactions between NtDD and CeDD surfaces and IDR (domain boundaries marked by dashed lines; color scale: number of contacts averaged over simulation time). Contacts within folded domains are omitted (set to zero) due to rigid-body constraints applied during the simulation. Note extensive contacts with C-terminal RGG motif denoted with red arrow. **(E)** Interdomain interactions drive IDR compaction in single-chain simulations. Main plot compares average intrachain distance, R_ij_, between the i^th^ and j^th^ residues of the isolated IDR (orange) and IDR in full-length (FL) context (blue). Attenuated R_ij_ expansion observed in FL context demonstrates interactions with the folded domains drive significant compaction of IDR. **Inset:** Violin plots of radius of gyration (R_g_) distributions further confirm restricted spatial dimensions of IDR in FL context compared to isolation. **(F)** Interaction networks in the condensed phase from CG slab simulations. Left contact map shows intermolecular contacts per chain, and right map shows intramolecular contacts per chain (domain boundaries marked by dashed lines; color scale: number of contacts averaged over simulation time). **(Center inset)** Representative 3D snapshot of simulated PGL-3 condensate illustrates a multi-chain, densely packed condensed phase.

To compare the dynamics of PGL-3 WT in the foci and cytoplasm, we performed FCS. FCS measures molecular diffusion based on fluorescence signal fluctuations across a range of lag times within a small confocal volume (**Fig. 2B**), which enables assessment of condensate dynamics *in vitro* and *in vivo*^51–53^. We performed FCS on foci in 25 anterior TRNs in day 2 adults by focusing the confocal volume inside the foci and computing the autocorrelation function (ACF) of the measured fluorescence fluctuations. The ACF curves reveal that the overall correlation decay time for PGL-3 WT in the foci was significantly longer (diffusion half-time, ∼720 ms) compared to PGL-3 WT in the cytoplasm (∼25 ms) (**Fig. 2C**). As protein diffusion in crowded or intracellular systems strongly deviates from simple diffusion models, we fit ACF traces to an anomalous diffusion model to calculate the apparent diffusion coefficient (D_app_)^54–57^. This analysis revealed that the longer decay time for PGL-3 WT in the foci corresponds to a nearly 20-fold decrease in the D_app_ of the PGL-3 molecules in the foci (D_app_ = 0.024 ± 0.01 µm^2^/s) compared to in the cytoplasm (D_app_ = 0.52 ± 0.25 µm^2^/s) (**Fig. 2C**). The D_app_ for PGL-3 WT in foci is of similar magnitude as values previously reported for dynamic liquid-like condensates^42,58^, and as expected, PGL-3 WT is far less dynamic within the condensate than in the cytoplasm. Taken together, *in vivo* FRAP and FCS data suggest that PGL-3 WT forms dynamic, liquid-like condensates when ectopically expressed in *C. elegans* TRNs, allowing us to test PGL-3 as a modifiable substrate to address how altering condensate properties might tune their localization as cargo into exophers.

### Inter-domain electrostatic interactions provide a mechanistic basis for PGL-3 phase separation

Having established that we can measure PGL-3 condensate dynamics in TRNs and observe PGL-3 WT in exophers, we next sought to engineer PGL-3 to generate a set of variants to assess how changes to condensate material properties might modulate exopher cargo recruitment. In designing new PGL-3 variants, we leveraged the current understanding of molecular grammar features of proteins that govern phase behavior and dynamics^48,59–62^.

Phase separation is often driven by multivalent weak intermolecular interactions, which may be between IDRs, or between folded domains and IDRs in multidomain proteins such as PGL-3^61^. Specifically, interdomain electrostatic interactions can modulate both phase behavior and material properties of condensates^63–66^. The PGL-3 primary sequence reveals an enrichment in charged residues, and electrostatic surface calculations show that both folded domains present acidic surfaces, while the RGG-rich region at the C-terminus of the IDR provides a complementary basic tail (**Fig. 2D**). This charge complementarity predicts extensive electrostatic interaction potential between the FD (NtDD and CeDD) and the IDR, which may play a central role in its phase behavior^64,65^.

To test this prediction, we performed single-chain simulations of the full-length (FL) PGL-3 protein. Contact analysis confirmed that the IDR tail forms extensive contacts with the folded domains, particularly among long-range residue pairs (upper region of the map, **Fig. 2D** right), which are typically contact-depleted due to chain-connectivity constraints. We assessed the interdomain interactions by comparing FL and IDR-only simulations. Intrachain distance scaling (R_ij_) and the radius-of-gyration (R_g_) distributions both show that the IDR is significantly more compact in the FL context than in isolation (**Fig. 2E**), confirming that FD:IDR contacts contribute to collapse of the IDR chain, driven by charge complementarity^64,67^.

To capture the interaction network in the condensed phase, we performed slab-geometry phase-coexistence simulations, which simultaneously capture inter- and intramolecular contacts (**Fig. 2F**). Consistent with prior dense-phase simulations, intrachain contacts reorganize into intermolecular interactions in the condensed phase, generating a more uniform and spatially homogenized contact network^68,69^. Nevertheless, the FD:IDR contact enrichment persists, and the same interaction hotspot between the acidic FD region and the basic RGG-rich region remains faintly detectable, particularly at the acidic patch located between NtDD and CeDD. This outcome confirms that the RGG region promotes FD:IDR electrostatic interactions, likely underpinning PGL-3 phase separation. These findings suggest a potential strategy for modulating PGL-3 phase behavior by altering the interdomain electrostatic interactions.

### The RGG-rich region in the PGL-3 IDR modulates condensate dynamics *in vitro*

We established in **Fig. 1C** that dimerization is essential for PGL-3 phase separation in TRNs. However, prior studies demonstrate that mutations in the IDR can modulate phase behavior and material properties of PGL-3 condensates by altering the number and strength of weak multivalent interactions^42,50^. Notably, arginine interactions are known to enhance phase separation of IDRs^59,60,70^, and our prior work has shown that Arg can form multiple favorable contacts with other residues by cation-pi and hydrogen bonding interactions^48^. The enhanced interaction propensity of Arg relative to Lys has been attributed not only to cation-pi interaction strength but also to differential dehydration between the two residues^69,71,72^. Furthermore, the increased interaction strength of Arg over Lys leads to a significant increase in the viscosity of condensates^48,73^. Arg and Lys residues in the PGL-3 IDR, especially the RGG region, provide a potential handle for tuning phase behavior and emergent material properties.

We designed two PGL-3 variants by mutating Arg (R) to Lys (K) or Lys to Arg within the PGL-3 IDR, keeping the folded domains (PGL-3_1-450_) unchanged. In the R→K variant, all 15 Arg are mutated to Lys (PGL-3_1-450_-IDR_R→K_), and in the K→R variant, all nine Lys residues are replaced with Arg (PGL-3_1-450_-IDR_K→R_). Both R→K and K→R variants maintain the same net charge as WT IDR, while differing in the number of Arg and Lys residues, which may alter interaction strength (**Fig. 3A**). For simplicity, we will refer to PGL-3_1-450_-IDR_R→K_ as PGL-3_R→K_ and PGL-3_1-450_-IDR_K→R_ as PGL-3_K→R_. To facilitate *in vitro* purification and characterization of the PGL-3 variants, proteins include a six-histidine and mNG tag at the PGL-3 N-terminus.

**Figure 3:**
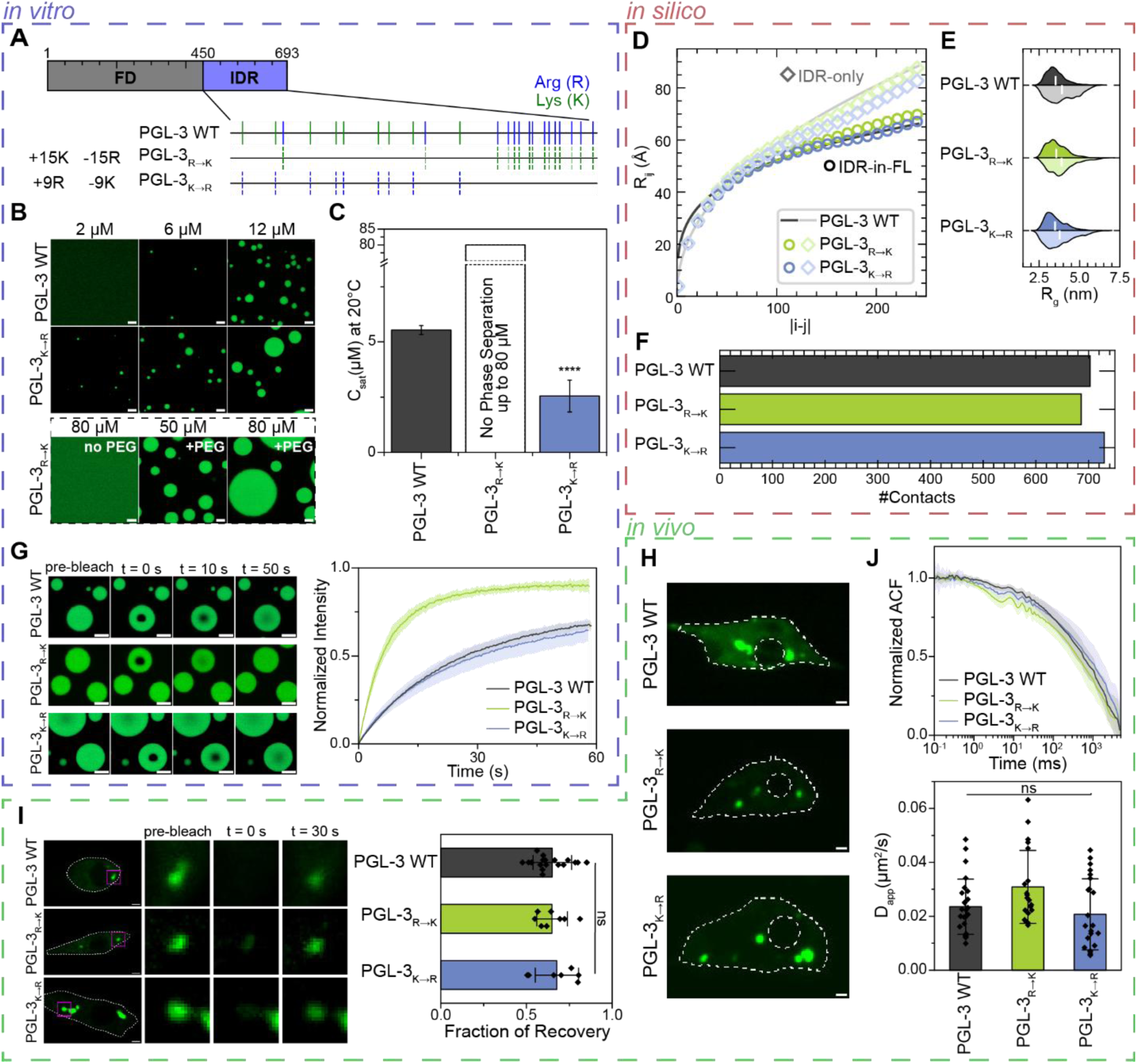
Tuning PGL-3 phase behavior through mutagenesis of the intrinsically disordered region. **(A)** Schematic showing mutations made in PGL-3_R→K_ and PGL-3_K→R_. Blue and green lines denote Arg and Lys residues, respectively, within WT IDR. Dashed lines represent R→K and K→R mutations. **(B)** Representative confocal microscopy images of purified proteins incubated at room temperature (∼20 °C) for 1 hour illustrating *in vitro* concentration-dependent phase separation of PGL-3 variants at 150 mM NaCl, 20 mM Tris, 1 mM DTT, pH 7.5. PGL-3_R→K_ does not phase separate in the absence of crowding agent up to 80 µM but does with 4% PEG 3500. Scale bar: 5 µm. **(C)** Phase separation quantified by measuring saturation concentration (C_sat_) at 20 °C through sedimentation assay. Data represents mean ± SD (n=6 from two independent experiments). Welch’s t-test (****p<0.00001). **(D)** R_ij_ profiles compare IDR in isolation (lighter) with IDR in FL context (darker) for PGL-3 WT (grey), PGL-3_R→K_ (green), and PGL-3_K→R_ (blue), calculated from CG single-chain simulations. All three variants show reduced R_ij_ scaling in FL context relative to IDR-only condition, indicating tethering to folded domains compacts the IDR. **(E)** Violin plots of R_g_ distributions of the IDR for each variant in IDR-only state (light) and in FL context (dark) supporting IDR compaction in full-length context. **(F)** Total number of contacts per chain in condensed phase from CG slab simulations. X-axis scale is the sum of all contacts across all residues across simulation time. **(G) (Left)** Representative confocal images for PGL-3 condensates from FRAP experiments pre-bleach, bleach (t=0 s), and post bleach (t=10 s, 50 s). Scale bars: 5 µm. **(Right)** Normalized FRAP recovery curves comparing condensates formed by PGL-3 variants. Note, *in vitro* experiments for PGL-3_R→K_ were conducted with 4% PEG 3500. Data represents mean ± S.D. (n=8-9 from at least two independent experiments). **(H)** Representative images of *C. elegans* ALM neurons expressing PGL-3_R→K_ (ZB5917) and PGL-3_K→R_ (ZB5918) (**Table S2**), showing foci formation *in vivo* (representative of n=14-24 images). Images oriented anterior left and posterior right. Soma and nucleus outlined in white; scale bars: 1 μm. **(I) (Left)** Representative FRAP images of anterior TRNs with soma outlined in white and bleached foci in magenta box. Zoom insets are pre-bleach, bleach, and 30 s post-bleach timepoints. Scale bars: 1 μm. **(Right)** Bar graph represents fraction of recovery based on recovery plateau post bleach. FRAP recovery traces found in **SI Fig. 4**. PGL-3 WT data same as in **Fig. 2A**. Data represents mean ± S.D. (n=7-19 individual foci from distinct neurons). Welch’s ANOVA and Games-Howell MCT. **(J) (Top)** Normalized FCS ACF traces from anterior TRNs. **(Bottom)** D_app_ extracted from ACF data fit to anomalous diffusion model. The extracted anomalous diffusion exponents (α) are shown in **SI Fig. 7**. PGL-3 WT data same as in **Fig. 2C** foci. Data represents mean ± S.D. (n=22-25 ACF traces). Welch’s ANOVA and Games-Howell MCT.

We reconstituted condensates formed by PGL-3 WT, PGL-3_R→K_, and PGL-3_K→R_ *in vitro* in physiological buffer (150 mM NaCl, 20 mM Tris, 1 mM DTT, pH 7.5) and determined their concentration-dependent phase behavior by fluorescence microscopy (**Fig. 3B**). We observed droplet formation for PGL-3 WT at ∼6 μM, whereas the PGL-3_K→R_ variant formed droplets at concentrations as low as ∼2 µM. In contrast, we found that the PGL-3_R→K_ variant did not phase separate even up to protein concentrations as high as 80 µM. Since molecular crowding agents such as PEG can be added to promote phase separation^74^, we introduced 4% PEG 3500 to our physiological buffer, resulting in PGL-3_R→K_ phase separating at ∼30 μM (**Fig. 3B**). We also performed a complementary sedimentation assay on the three constructs to determine the saturation concentration (C_sat_), which identifies the minimum concentration at which the solution separates into dense and dilute phases at the experimental temperature (20 °C) (**Fig. 3C**). The C_sat_ measurements by sedimentation assay were consistent with the microscopy observations (**Fig. 3B, C**). As expected, PGL-3_K→R_ showed increased phase separation propensity while PGL-3_R→K_ showed diminished phase separation propensity relative to PGL-3 WT^48,59^ .

To identify how these sequence changes alter the interaction profile, we performed single-chain simulations of PGL-3_R→K_ and PGL-3_K→R_ for comparison to PGL-3 WT. Contact analysis revealed that the IDR of PGL-3_R→K_ forms fewer contacts in both the FL and IDR-only contexts, whereas the IDR of PGL-3_K→R_ forms more contacts (**SI Fig. 2A**), consistent with the established roles of Arg and Lys in LLPS^59^. These differences are directly reflected in chain conformation. R_ij_ and R_g_ analyses highlight the influence of FD:IDR interdomain interactions, as all variants adopt more compact conformations in the FL context relative to their isolated IDR counterparts (**Fig. 3D, 3E**). Among them, PGL-3_R→K_ exhibits a slightly more expanded IDR ensemble, consistent with its reduced contacts, while PGL-3_K→R_ shows the most collapsed IDR conformation in the IDR-only construct. Notably, the IDR of PGL-3_K→R_ converges toward WT-like compaction in the FL context, likely because the folded domains constrain the IDR by engaging in FD:IDR contacts, limiting further IDR collapse.

The same interaction hierarchy persists in the condensed phase (**SI Fig. 2B**). Contact analysis of slab-geometry phase-coexistence simulations reveals that PGL-3_K→R_ exhibits the highest total number of contacts and PGL-3_R→K_ the lowest, with PGL-3 WT intermediate (**Fig. 3F**), mirroring the trends observed in the single-chain simulations. These results underscore the enhanced interaction capacity introduced by additional Arg residues in the IDR. These computational findings provide a mechanistic explanation for the observed *in vitro* LLPS propensity of the R/K variants, linking sequence-encoded interaction tuning to both single-chain behavior and dense-phase contact networks.

We next assessed the *in vitro* dynamics of PGL-3 WT, PGL-3_K→R_, and PGL-3_R→K_ condensates using FRAP (**Fig. 3G**). We photobleached a small region within the condensates and observed the fluorescence recovery as unbleached fluorescent molecules diffused into the bleached region. We found that PGL-3 WT and PGL-3_K→R_ condensates exhibited similar fluorescence recovery. By contrast, PGL-3_R→K_ condensates (formed in the presence of 4% PEG) exhibited more rapid and near-full fluorescence recovery, suggesting enhanced internal dynamics as compared to PGL-3 WT and PGL-3_K→R_. We calculated the apparent diffusion coefficient (D_app_) by fitting the infinite boundary model to the FRAP traces^75^ (Methods, **SI Fig. 3**). The D_app_ for PGL-3 WT and PGL-3_K→R_ condensates are similar in magnitude with ∼0.01 µm^2^/s, whereas for the PGL-3_R→K_ variant, we found the D_app_ to be ∼0.04 µm^2^/s. Thus, R→K mutations within the PGL-3 IDR reduce interaction strength and increase C_sat_, as well as make condensates more dynamic. However, though K→R mutations increase interaction strength and decrease C_sat_, they do not significantly alter condensate dynamics relative to WT. Together, these data support that interactions mediated by the RGG-rich region are important contributors to PGL-3 phase separation and dynamics.

### PGL-3 K/R variants phase separate with liquid-like properties *in vivo*

Our core question is whether PGL-3 variants expressed *in vivo* form foci with different material properties and are differentially included as exopher cargo. To address this question, we next sought to compare the dynamics of PGL-3 WT, PGL-3_R→K_, and PGL-3_K→R_ in *C. elegans* TRNs. We expressed mNG-tagged PGL-3_R→K_ (strain ZB5917) and mNG-tagged PGL-3_K→R_ (strain ZB5918) using the *mec-7* promoter to drive TRN expression with the TRN mCherry allele (**Table S2**). We found that PGL-3_R→K_ and PGL-3_K→R_ both formed foci in the ALM neurons of day 2 adults, similar to PGL-3 WT, with some variation in size and number of foci (**Fig. 3H, SI Fig. 1**). (In the expression lines we generated, each PGL-3 variant is integrated as a single-copy insertion to maintain similar expression levels. We note, however, that the chromosomal insertion site, which can modulate expression levels, is random; expression levels measured by fluorescence for these strains are shown in **SI Fig. 1**).

To assess *in vivo* dynamics of PGL-3 variants, we performed FRAP experiments on foci in anterior TRNs of day 2 adults (**Fig. 3I**). FRAP analysis suggests a 60-70% fraction of recovery within 30 s for PGL-3 WT, PGL-3_R→K_ and PGL-3_K→R_, reflecting the dynamic nature of the PGL-3 foci *in vivo* (**Fig. 3I, SI Fig. 4**). To more accurately quantify changes in the protein diffusion within the foci, we performed FCS experiments on foci in anterior TRNs of day 2 adults. The ACF traces show that PGL-3 WT, PGL-3_R→K_, and PGL-3_K→R_ have similar decay curves (**Fig. 3J**). The calculated D_app_ for PGL-3 WT and PGL-3_K→R_ foci in the soma are similar (0.024 ± 0.01 and 0.021 ± 0.013 µm^2^/s, respectively) and slightly greater for PGL-3_R→K_, 0.031 ± 0.014 µm^2^/s (**Fig. 3J**).

Overall, our results indicate that the PGL-3 K/R variants form dynamic condensates *in vivo* with the differences between D_app_ for the variants largely reduced relative to those observed *in vitro*. The resulting limited dynamics range of PGL-3 K/R variants *in vivo* prompted us to design additional PGL-3 variants to expand that range as we sought to answer whether material properties govern exopher cargo selection.

### Artificial intrinsically disordered polypeptides provide a platform to further engineer PGL-3 condensates with tunable physical properties

To generate PGL-3 condensates with a wider range of material properties for exopher cargo studies, we expanded our approach by swapping the native PGL-3 IDR with artificial intrinsically disordered polypeptides (AIDPs). AIDPs are repeat sequences that are inspired by naturally occurring IDRs^48,76^. We and others have previously shown that these AIDP sequences allow programmability of homotypic (AIDP-AIDP) interactions with effective control of phase separation propensity and condensate material properties *in vitro* and *in vivo*^48,76,77^. This previous work prompted us to ask whether AIDPs might be utilized to tune the multivalent interactions that drive phase separation in proteins with multidomain architecture. Although myriad options exist for AIDPs, two stood out as rational choices based on our previous study^48^: 1) (GRGDSPYS)_25_, twenty-five repeats of an eight amino acid zwitterionic sequence, which features several residues and interactions believed to play key roles in modulating phase separation—we call this sequence here AIDP; and 2) (GRG<u>N</u>SPYS)_25_, a cationic variant referred to here as AIDP+, selected to complement the net negative surface charge of the PGL-3 folded domains. We therefore swapped the native PGL-3 IDR (451-693) with the AIDP and AIDP+ sequences, while maintaining the folded domains (1-450) unchanged (**Fig. 4A**).

**Figure 4:**
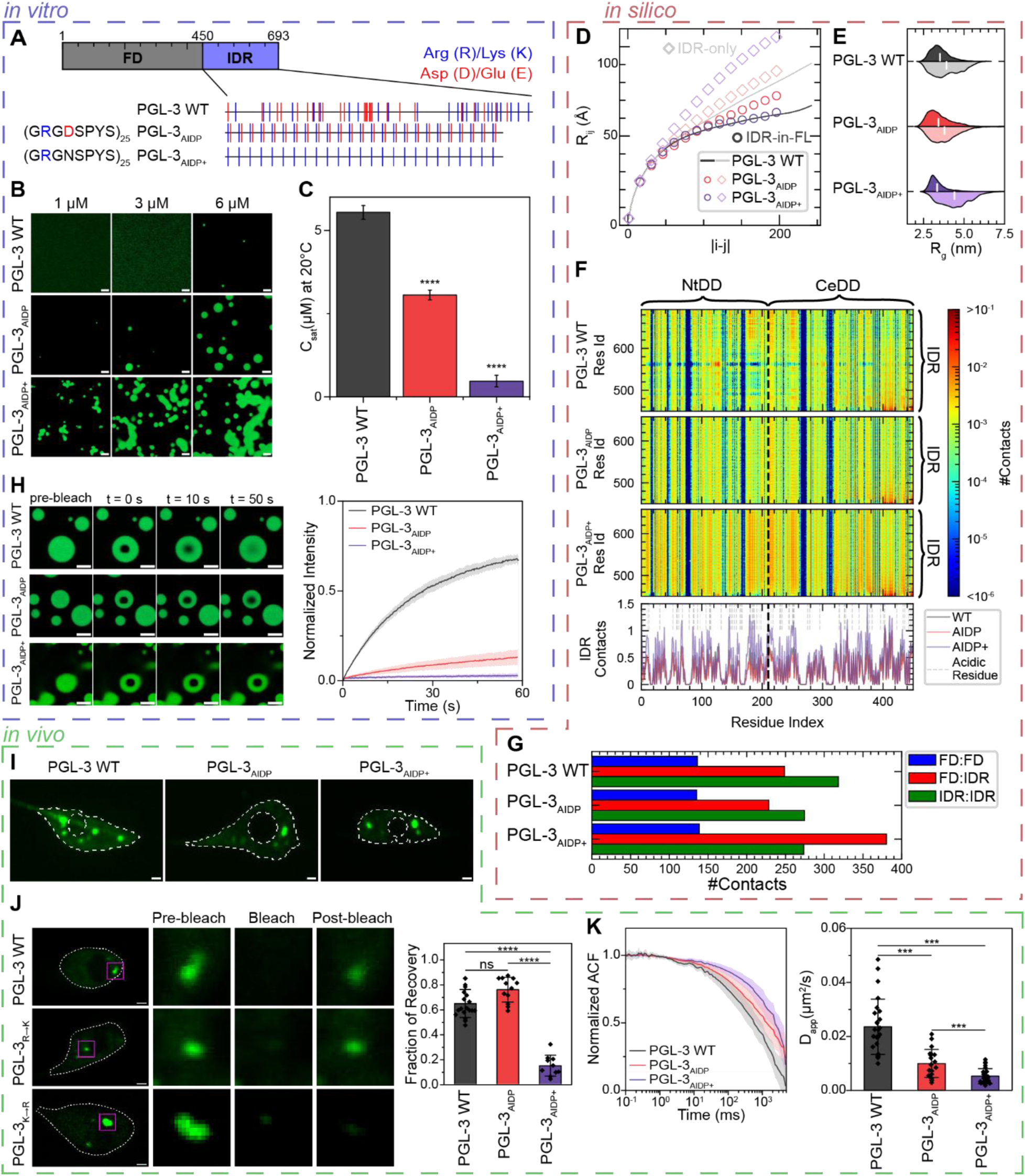
PGL-3 AIDP chimeras display altered phase behavior and condensate dynamics. **(A)** Schematic showing PGL-3 IDR replacement with AIDP and AIDP+. Blue lines denote basic residues Arg and Lys. Red lines denote acidic residues Asp and Glu. Note, PGL-3_AIDP_ and PGL-3_AIDP+_ have linker sequences, RSEFNG and RSEFDP, respectively, between the FD and IDR sequences (**Table S1**). **(B)** Representative confocal microscopy images of purified proteins incubated at room temperature (∼20 °C) for 1 hour illustrating *in vitro* concentration-dependent phase separation of PGL-3 WT, PGL-3_AIDP_, and PGL-3_AIDP+_ at 150 mM NaCl, 20 mM Tris, 1 mM DTT, pH 7.5. Scale bar: 5 µm. **(C)** Phase separation quantified by measuring C_sat_ at 20 °C through sedimentation assay. PGL-3 WT data same as in **Fig. 3C**. Data represents mean ± SD (n=6 for all variants from two independent experiments). **(D)** R_ij_ profiles from single-chain simulations comparing isolated IDR (lighter) with the IDR in the FL context (darker) for PGL-3_AIDP_ (red) and PGL-3_AIDP+_ (purple). PGL-3 WT profile (grey) shown for reference. PGL-3_AIDP_ shows modest compaction relative to its IDR-only baseline. PGL-3_AIDP+_ undergoes pronounced compaction in FL context. **(E)** Violin plots of R_g_ distributions of the IDR for each variant in the IDR-only state (lighter) and in the FL context (darker), highlighting compaction of PGL-3_AIDP+_ in FL context. **(F)** Two-dimensional FD:IDR interdomain contact maps (FD encompasses NtDD and CeDD) in CG slab simulations (color scale: number of contacts averaged over simulation time). **(Below)** Cumulative number of FD:IDR contacts at each FD residue across simulation time. PGL-3_AIDP+_ establishes substantially stronger and broadly distributed FD:IDR contacts, contrasting with the minimal FD:IDR engagement of PGL-3_AIDP_. **(G)** Molecular contacts in CG slab simulations by domain pair. X-axis scale is the sum of all contacts per domain pair across simulation time. Contact counts reveal PGL-3_AIDP+_ exhibits dramatic increase in FD:IDR contacts relative to WT. PGL-3_AIDP_ displays reduced FD:IDR contacts, shifting toward an IDR:IDR-driven network. **(H) (Left)** Representative confocal images for PGL-3 condensates from FRAP experiments pre-bleach, bleach (t=0 s), and post bleach (t=10 s, 50 s); scale bars: 5 µm. **(Right)** Normalized FRAP recovery curves showing change in dynamics of the condensates formed by PGL-3 variants. PGL-3 WT data same as in **Fig. 3G**. Data represents mean ± SD (n=8-9 for all variants from at least two independent experiments). **(I)** Representative images of *C. elegans* ALM neurons expressing PGL-3_AIDP_ (ZB5919) and PGL-3_AIDP+_ (ZB5920) (**Table S2**), showing foci formation *in vivo* (representative of n=20-26 images). Images oriented anterior left and posterior right. Soma and nucleus outlined in white; scale bars: 1 μm. **(J) (Left)** Representative FRAP images of anterior TRNs with soma outlined in white and bleached foci in magenta box. Zoom insets are pre-bleach, bleach, and 30 s post-bleach timepoints. Scale bars: 1 μm. **(Right)** Bar graph representing fraction of recovery based on recovery plateau post bleach. FRAP recovery traces found in **SI Fig. 4**. PGL-3 WT data is the same as in **Fig. 2A**. Data represents mean ± S.D. (n=10-19 individual foci from distinct neurons). Welch’s ANOVA and Games-Howell MCT (****p < 0.0001). **(K) (Left)** Normalized FCS ACF traces from anterior TRNs. **(Bottom)** D_app_ extracted from ACF data fit to anomalous diffusion model. The extracted anomalous diffusion exponents (α) are shown in **SI Fig. 7**. FCS analysis shows PGL-3_AIDP+_ has the slowest diffusion followed by PGL-3_AIDP_. PGL-3 WT data same as in **Fig. 2C** foci. Data represents mean ± S.D. (n=21-25 individual FCS traces). Welch’s ANOVA and Games-Howell MCT (*** p < 0.001; ****p < 0.0001).

We expressed and purified the two PGL-3 AIDP chimeric proteins (PGL-3_1-450_-AIDP and PGL-3_1-450_-AIDP+), referred to in this study as PGL-3_AIDP_ and PGL-3_AIDP+_, respectively. Similar to the PGL-3 K/R variants, PGL-3_AIDP_ and PGL-3_AIDP+_ demonstrated protein-concentration-dependent phase behavior in physiological buffer (150 mM NaCl, 20 mM Tris, 1 mM DTT, pH 7.5), confirmed by fluorescence microscopy (**Fig. 4B**). We found the onset of droplet formation at protein concentrations as low as 2 µM for PGL-3_AIDP_, and less than 1 µM for PGL-3_AIDP+_ (**Fig. 4B**). We independently measured the C_sat_ by sedimentation assay and obtained results consistent with our microscopy: C_sat_ for PGL-3_AIDP_ and PGL-3_AIDP+_ were ∼2 µM and ∼0.7 µM, respectively, as compared to ∼6 µM for PGL-3 WT (**Fig. 4C**). These results show that AIDPs can enhance the phase separation propensity of PGL-3 and provide another handle to modulate condensate physical properties.

Single-chain simulations show that loss of the basic (RGG) tail in PGL-3_AIDP_ weakens FD:IDR interactions (**SI Fig. 5**), leaving the IDR chain more expanded in the FL context compared with PGL-3 WT (**Fig. 4D**). PGL-3_AIDP+_, conversely, forms extensive FD:IDR contacts in the FL context due to electrostatic complementation (**SI Fig. 5**), driving marked chain compaction as evidenced by both its R_ij_ scaling and R_g_ values (**Fig. 4D, 4E**). In the isolated IDR analysis, this trend reverses: AIDP+ adopts a swollen, highly expanded ensemble consistent with charge-repulsion-driven extension and minimal intradomain contacts, whereas AIDP exhibits stronger IDR:IDR intradomain contacts that produce a more collapsed conformation (**SI Fig. 5**). Nevertheless, FD:IDR interactions in the FL context consistently shift the IDR toward more compact conformations relative to the IDR-only constructs across all variants.

To further examine how these interaction networks shape phase behavior, we performed slab-geometry phase-coexistence simulations. Within this common framework, domain-pair contact analysis recapitulates differences observed in the single-chain interactions: PGL-3_AIDP+_ is enriched in FD:IDR contacts, whereas PGL-3 WT and PGL-3_AIDP_ show a greater relative contribution from IDR:IDR contacts than from FD:IDR contacts (**Fig. 4F, 4G, SI Fig. 5**). The 2D contact maps resolve the residue-level basis of these differences. Relative to PGL-3 WT, the IDR of PGL-3_AIDP+_ makes broadly distributed contacts with the FD, with pronounced enrichment near acidic FD residues, whereas FD:IDR contacts are depleted in PGL-3_AIDP_ (**Fig. 4F**). These contact maps point to enhanced electrostatic complementarity between the PGL-3_AIDP+_ IDR and the FD as a key feature of its distinct interaction network and altered phase behavior.

Next, we assessed the dynamic properties of the PGL-3 AIDP chimeras *in vitro*. For PGL-3_AIDP_ condensates, the spherical droplet morphology suggests that PGL-3_AIDP_ condensates are liquid-like. In contrast, PGL-3_AIDP+_ condensates showed amorphous morphology arising from incomplete droplet fusion events, suggesting a gel-like or dynamically arrested state (**Fig. 4B**). FRAP analysis showed a significantly slower recovery rate for PGL-3_AIDP_ condensates (20% recovery in 60 s) compared to PGL-3 WT (70% in 60 s), indicating reduced dynamics of PGL-3_AIDP_ condensates (**Fig. 4H**). In contrast, PGL-3_AIDP+_ condensates showed no observable fluorescence recovery, suggesting the condensed proteins are immobile over time scales of at least 1 min (**Fig. 4H**). Overall, the two PGL-3 AIDP chimeras showed drastically slower *in vitro* dynamics when compared to PGL-3 WT and R/K variants, highlighting the application of artificial repeat sequences in tuning physical properties of the condensates.

### PGL-3_AIDP_ and PGL-3_AIDP+_ form condensates in *C. elegans* TRNs, consistent with *in vitro* dynamics measurements

Having established greatly reduced dynamics *in vitro* for PGL-3_AIDP_ and PGL-3_AIDP+_ variants, we next sought to assess their dynamics when expressed in TRNs, so that we could ultimately address whether condensate properties affect exopher cargo recruitment. We expressed PGL-3_AIDP_ (strain ZB5919) and PGL-3_AIDP+_ (strain ZB5920) in TRNs using the same N-terminal mNG tag and *mec-7* promoter that we used for PGL-3 WT expression, along with the TRN mCherry allele (**Fig. 4I**) (**Table S2**). We found that PGL-3_AIDP_ and PGL-3_AIDP+_ formed foci in ALM neurons of day 2 adults, similar to PGL-3 WT and R/K variants, with some variation in size and number (**Fig. 4I, SI Fig. 1**).

We then used FRAP and FCS to examine the dynamics of PGL-3_AIDP_ and PGL-3_AIDP+_ foci *in vivo* in anterior TRNs. We found that PGL-3_AIDP_ foci have 60%-70% fluorescence recovery in 30 s, suggesting liquid-like dynamics (**Fig. 4J**). Although *in vitro* PGL-3_AIDP_ droplets exhibited slower FRAP recovery than PGL-3 WT, *in vivo* foci showed relatively quick recovery, suggesting fast exchange with the cytoplasm and/or a weakened protein-interaction network within the condensates *in vivo*. In contrast, PGL-3_AIDP+_ foci *in vivo* showed less than 20% FRAP recovery in 30 s, which is markedly slower than the other PGL-3 variants (**Fig. 4J**) and consistent with *in vitro* measurements.

To measure protein diffusion within the foci, we performed FCS on foci in anterior TRNs of day 2 adults, comparing PGL-3 WT, PGL-3_AIDP_, and PGL-3_AIDP+_. Our analysis showed that ACF traces for PGL-3_AIDP+_ exhibit slower decay than PGL-3_AIDP_, which exhibits slower decay than PGL-3 WT, indicating that PGL-3_AIDP_ and PGL-3_AIDP+_ experience slower diffusion within condensates compared to PGL-3 WT (**Fig. 4K**). The calculated D_app_ is 0.024 ± 0.01 µm^2^/s for PGL-3 WT, 0.01 ± 0.005 µm^2^/s for PGL-3_AIDP_, and 0.005 ± 0.003 µm^2^/s for PGL-3_AIDP+_ (**Fig. 4K**). Interestingly, the measured D_app_ for PGL-3_AIDP+_ is of the same order of magnitude as the *in vivo* D_app_ for MEG-3, which is known to form a gel-like sub-phase of P granules^42,58^. Overall, FCS indicates that the internal diffusion of molecules within the foci *in vivo* parallels the trends from *in vitro* FRAP, although differences are reduced *in vivo* compared to *in vitro*. Importantly, we showed that we can modulate the dynamics of a naturally occurring protein *in* TRNs by substituting AIDPs for the native IDR. Our findings also establish PGL-3_AIDP_ and PGL-3_AIDP+_ as suitable substrates to study exopher sorting of condensates with slower dynamics than PGL-3 WT.

### Engineered PGL-3 condensates exhibit a spectrum of viscoelastic properties *in vitro*

Thus far, we have established that the PGL-3 variants form condensates *in vitro* and *in vivo* with different dynamics. However, to differentiate condensate material properties (i.e., liquid-like versus gel-like) and complement diffusion dynamics of PGL-3 variants, we directly probed condensate viscoelasticity. An emerging body of work suggests that protein phase separation gives rise to condensates with reversible network-like structures that lead to viscoelasticity, a soft-matter property that contributes to condensate function or dysfunction in living cells^42,48,78–82^. Viscoelasticity plays a key role in determining how condensates respond to the mechanically active intracellular environment, which can influence cargo trafficking^83,84^.

Here, we quantified the viscoelasticity of the condensates *in vitro* using video particle tracking (VPT) passive microrheology^48,73^. Briefly, this method measures the rheological properties of the local environment by tracking the Brownian motion of tracer beads embedded within the condensates. The mean squared displacements (MSDs) of the tracer beads can be used to calculate the viscoelastic behavior of the condensates. The slope of the MSD vs. lag time curve on a log-log plot corresponds to the diffusivity exponent (α) that can be used to differentiate between purely viscous fluids (α ∼ 1) and viscoelastic materials (0 < α < 1). We mixed fluorescent tracer beads with the protein samples *in vitro* (see Methods) and recorded the trajectories of beads that partitioned within the condensates (**Fig. 5A**). The ensemble-averaged MSDs of beads in PGL-3 WT, PGL-3_K→R_, PGL-3_R→K_, and PGL-3_AIDP_ condensates increased linearly with lag-time, with α ∼ 1, suggesting a viscous, liquid-like environment (**Fig. 5B**). However, the observed MSD for PGL-3_AIDP+_ condensates showed sub-diffusive behavior (α ∼ 0.22) over the 1 s timescale probed, suggesting highly constrained motion of embedded tracer beads, also shown by representative bead trajectories (**Fig. 5A**).

**Figure 5:**
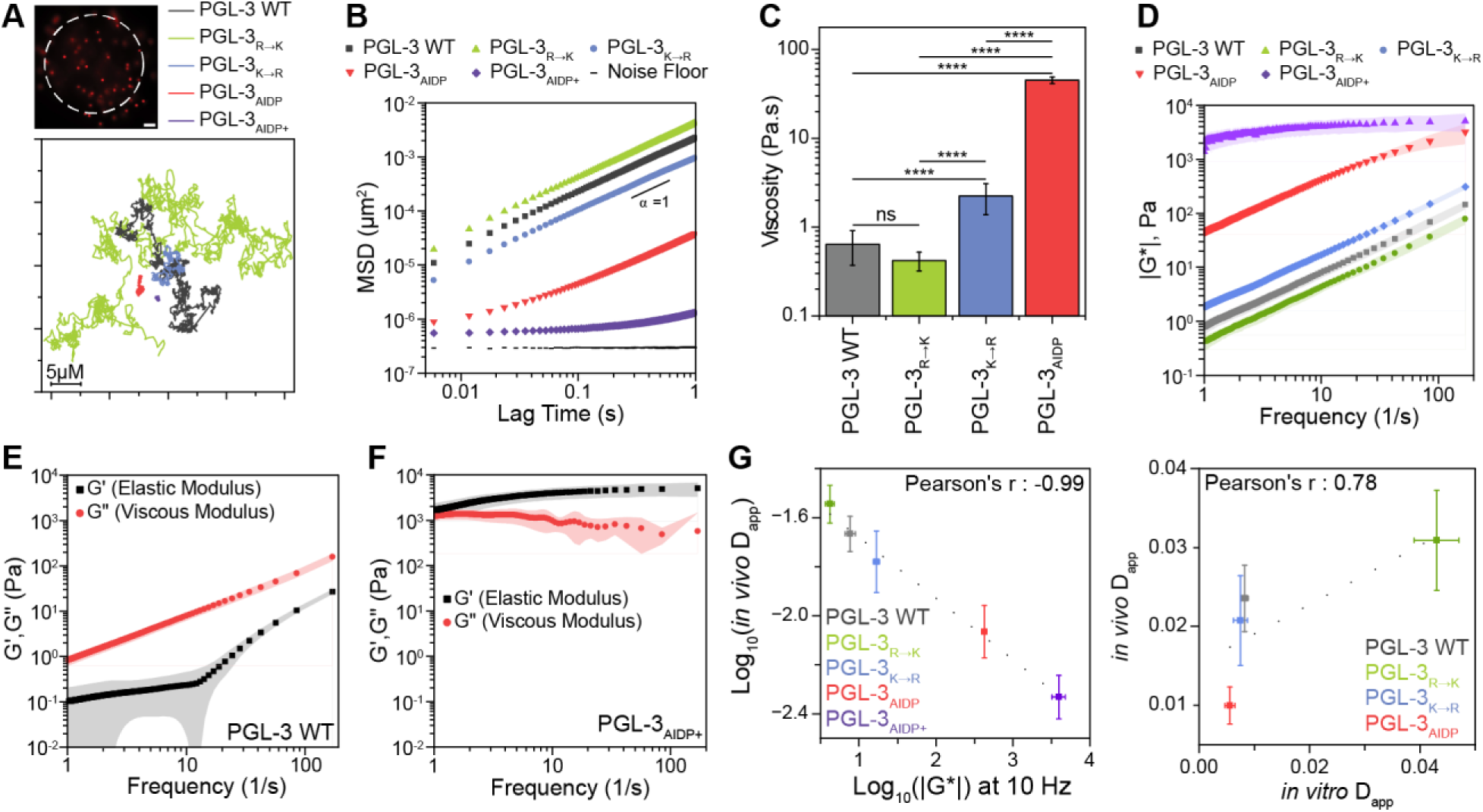
AIDPs alter mechanical properties of native protein condensates. **(A) (Top)** Representative image of 1 µm red fluorescent polystyrene tracer beads embedded in a PGL-3 WT droplet (outlined in white) for particle tracking microrheology. Scale bar: 5 µm. **(Bottom)** Representative particle trajectories over 1 s showing significant change in Brownian motion of beads within condensates for PGL-3 variants, as indicated. **(B)** Ensemble-averaged mean squared displacements (MSDs) of tracer beads within condensates at ∼1 hour after initiating phase separation. PGL-3_AIDP+_ condensates show sub-diffusive motion, whereas all other variants show normal diffusive behavior confirmed by diffusivity exponent α ∼ 1, based on fits to the entire MSD vs. lag time plots (n=8-9 different videos from two independent experiments for all variants). **(C)** *In vitro* viscosity calculated from MSDs using Stokes-Einstein relation. Due to deviation from normal diffusion (α < 1), viscosity was not determined for PGL-3_AIDP+_ condensates. Data represents mean ± S.D.; n same as in (B). Welch’s ANOVA and Games-Howell MCT (****p< 0.0001). **(D)** Magnitudes of the complex modulus (|G*|, where G*=G’+iG”) for five variants calculated from the algebraic conversion of MSDs in (B), illustrating differences in variant-specific condensate stiffness. Higher |G*| corresponds to greater resistance to deformation. **(E; F)** Measured frequency-dependent viscoelastic moduli (G’ = elastic modulus, G” = viscous modulus) of PGL-3 WT and PGL-3_AIDP+_ condensates, calculated from MSDs in (B). PGL-3 WT exhibits dominant viscous behavior with G” > G’, while PGL-3_AIDP+_ exhibits gel-like behavior with G’ > G” across all measured frequencies. **(G)** Plots showing strong correlation between mean *in vitro* condensate |G*| and *in vivo* condensate dynamics D_app_ from FCS (**Fig. 3J and 4K**), and between *in vivo* D_app_ from FCS and *in vitro* D_app_ from FRAP (**SI Fig. 3**). Error bars represent 95% confidence intervals for the mean values.

We next used the Stokes-Einstein equation to calculate the viscosity of the PGL-3 WT, PGL-3_K→R_, PGL-3_R→K_, and PGL-3_AIDP_ condensates from their MSDs. The viscosity measured for PGL-3 WT condensates was 0.64 ± 0.27 Pa·s (**Fig. 5C**). PGL-3_K→R_ showed a significant ∼3x increase in viscosity (2.23 ± 0.85 Pa·s), whereas PGL-3_R→K_ (with 4% PEG) condensates showed a slightly lower measured viscosity (0.42 ± 0.1 Pa·s) compared to WT. In contrast, PGL-3_AIDP_ condensates have a significantly higher viscosity of 45.12 ± 3.74 Pa·s. The α ∼ 1 indicates that PGL-3_AIDP_ condensates are predominantly liquid-like in nature, even with their higher viscosity. The high viscosity may explain the relatively slow FRAP recovery observed for PGL-3_AIDP_ condensates *in vitro* shown in **Fig. 4H**.

For further rheological assessment, we examined the frequency-dependent viscoelastic properties of the condensates to distinguish the timescales at which elastic or viscous mechanical behavior dominates. We first calculated the magnitude of the complex modulus (|G*(ω)|), a measure of a material’s overall resistance to deformation, for the PGL-3 variants from the MSDs using the Generalized Stokes-Einstein Relationship (GSER)^85^ (**Fig. 5D**). PGL-3_AIDP+_ exhibited the highest value over the entire frequency range tested. Across all variants, |G*| varied by three orders of magnitude, with amplitudes ranging from 4 Pa to 4 kPa at a frequency of ω = 10 Hz. We next estimated the viscous (G”) and elastic (G’) moduli as well as the loss tangent (tan δ = G”/G’), which identifies the timescales at which the condensates are viscoelastic fluids (tan δ > 1) versus viscoelastic solids (tan δ < 1), revealing a pronounced material contrast between the variants (**Fig. 5E, 5F, SI Fig. 6**). We found that PGL-3 WT condensates show G” > G’ across all probed frequencies, indicating predominantly viscous mechanical behavior (**Fig. 5E**). In contrast, PGL-3_AIDP+_ shows G’ > G” and a significantly increased G’ across all probed frequencies, indicating increased network elasticity and gel-like mechanical behavior (**Fig. 5F**). We attribute the rheology of PGL-3_AIDP+_ to a stable elastic network formed by strong physical crosslinks—reversible, non-covalent connections—resulting from strong inter-domain interactions between the PGL-3 folded domains and AIDP+, consistent with our simulations. The other variants maintain a predominantly viscous response, indicating the absence of a crosslinked network structure with an increase in condensate viscoelasticity in the order of PGL-3_AIDP_ > PGL-3_K→R_ > PGL-3 WT ≈ PGL-3_R→K_ (**SI Fig. 6**). The results indicate an important role of sequence-encoded interactions in strengthening the intra-condensate network, thereby modulating viscoelasticity.

An important aspect of condensate biophysics is assessing how well *in vitro* and *in vivo* biophysical measurements align with one another. We found a strong correlation between *in vitro* FRAP and *in vivo* FCS diffusion coefficients (Pearson’s r = 0.78) and a very strong correlation between logarithms of *in vivo* FCS diffusion coefficients and *in vitro* VPT viscoelastic moduli (Pearson’s r = 0.99) (**Fig. 5G**). This suggests that *in vitro* material properties provide a reliable predictor of *in vivo* diffusivity, although deviations from perfect correlation likely reflect additional *in vivo* interactions within the complex cellular milieu compared to the relatively simple buffers *in vitro*.

### Condensate material properties tune exopher cargo recruitment

Having characterized the material properties of a panel of PGL-3 variants *in vitro* and *in vivo*, we returned to the central question of this paper: whether condensate material properties influence exopher cargo sorting. First, we addressed whether the PGL-3 variants could be extruded by neurons into exophers at all. We imaged exopher events from the ALM neurons in day 2 adults fasted for 3 hours to promote exopher formation. We used the TRN mCherry allele (*bzIs166*) in the strains to enable us to identify both soma and exopher, with the soma distinguished by its intact axon. We found that for each strain, some or all of the mNG-tagged PGL-3 variant foci could be observed in the exopher compartment (**Fig. 6A**). Thus, the tested modifications to the IDR sequence do not prevent neurons from extruding PGL-3 proteins into exopher vesicles.

**Figure 6:**
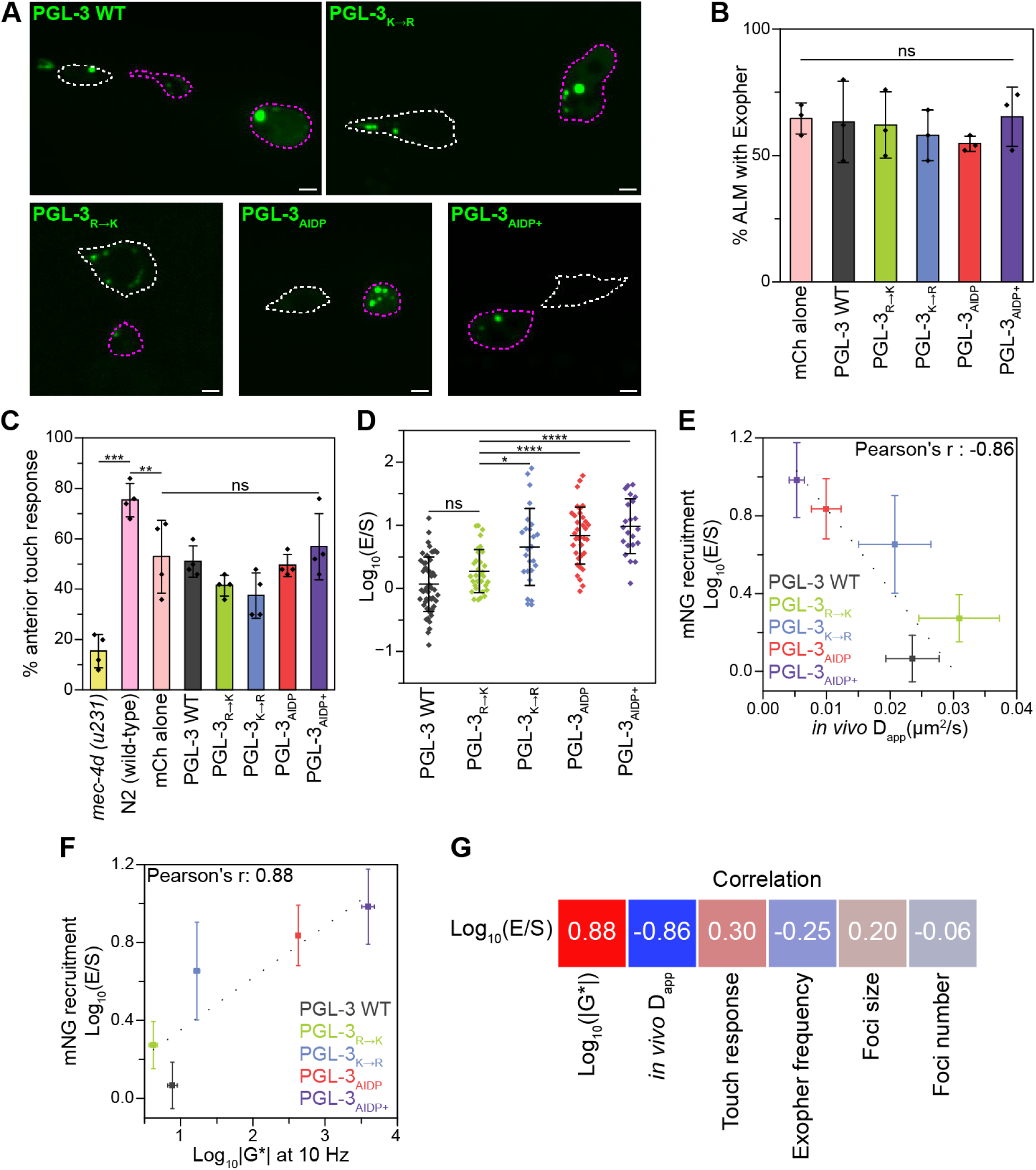
Exopher Formation and Recruitment Correlate with Condensate Dynamics in TRNs. **(A)** Representative images of exopher events in ALM neurons expressing PGL-3 variants. Images oriented anterior left and posterior right. Day 2 adults fasted for 3 hours prior to imaging; soma outlined in white; exophers outlined in magenta; scale bars: 2 µm. **(B)** Percent of day 2 adults fasted for 3 hours with an exopher from ALM neurons. Data represents mean ± S.D. (n=3 replicates of 50 animals counted). Cochran–Mantel–Haenszel test comparing each strain with PGL-3 WT. **(C)** Percent of day 1 adults that respond to anterior gentle touch assay. *mec-4d (u231)* = loss of all six TRNs, N2 = wild-type lab strain, mCh alone = bzIs166[*Pmec-4::mCherry*], PGL-3 variants are expressed with *bzIs16*6. Data represents mean ± S.D. (n=4 replicates of 10 animals tested 5 times each). One-way ANOVA and Dunnett’s MCT comparing each strain with PGL-3 WT (**p<0.01; ***p<0.001). **(D)** Localization of PGL-3 variants in soma and exopher. PGL-3 WT data repeated from **Fig. 1E**. Data represents mean ± S.D. (n=22-51 images per PGL-3 variant). One-way ANOVA and Tukey’s MCT (****p<0.0001). **(E)** Scatter plot representing correlation between exopher recruitment and *in vivo* condensate dynamics. Y-axis represents Log_10_(E/S) from (D). X-axis represents D_app_ calculated from FCS data in **Fig. 3J and 4K**. Dotted line represents negative linear relationship with Pearson’s r = -0.86. Error bars represent 95% confidence intervals for mean values. **(F)** Scatter plot representing correlation between exopher recruitment and *in vitro* condensate material properties. Y-axis represents Log_10_(E/S) from (D). X-axis represents complex modulus at 10 Hz calculated in **Fig. 5D**. Dotted line represents linear relationship with Pearson’s r = 0.88. Error bars represent 95% confidence intervals for mean values. **(G)** Heatmap of Pearson correlation coefficients calculated between Log_10_(E/S) and parameters measured in this study. Red indicates a strong positive correlation, and blue indicates a strong negative correlation. Of note, Log_10_(E/S) or exopher recruitment only correlates strongly with *in vivo* condensate dynamics and *in vitro* condensate material properties and not with other measured phenotypes

We next asked if expression of any of the PGL-3 variants could alter neuronal exopher production, motivated by our previous work demonstrating that the level of proteostress correlates with the relative frequency of exopher production^9^. Using the mCherry marker to identify exophers, we measured exopher levels in day 2 adults fasted for 3 hours. For all tested strains, introduction of the PGL-3 variants did not significantly alter exopher production compared to mCherry alone (**Fig. 6B**)—unlike Htt-polyQ128 aggregates, which stimulate exopher production^5^. This suggests that even the dynamically arrested PGL-3_AIDP+_ does not trigger impactful proteostatic stress in the neuron.

To evaluate functional neuronal health more directly, we assayed touch sensitivity in animals expressing the PGL-3 variants to measure the health and function of the anterior touch response circuit controlled by the TRNs^5,7,86^. Gently stroking the animal across the anterior portion of the body elicits a touch avoidance response in wild–type animals. Across 4 trials, we probed 40 animals for each PGL-3 variant, with 5 touches per animal. We found that, on average, the N2 wild-type animals (control lab strain) responded to 75.5% of touches, while the mutant *mec-4d(u231)*, in which the TRNs degenerate^87^, only responded to 15.5% of touches (**Fig. 6C**). The strain expressing mCherry in the TRNs responded to 53% of touches, and we found that expression of PGL-3 variants had no significant effect on touch sensitivity compared to mCherry alone (**Fig. 6C**). These data suggest that expression of the PGL-3 variants does not adversely affect neuronal health beyond the proteostress induced by mCherry expression. While exophers remove deleterious cargo^5,6,8^, these findings suggest that any changes in PGL-3 variant recruitment to exophers are not a response to proteotoxicity.

Finally, we asked whether particular PGL-3 variants might be differentially localized to exophers. Using the Log_10_(E/S) measure of relative exopher/soma distribution defined in **Fig. 1E**, we quantified and compared the distribution of mNG fluorescence for each PGL-3 variant. Relative to PGL-3 WT, for which Log_10_(E/S) = 0.067 ± 0.43, the other PGL-3 variants increasingly favor exopher compartment distribution. The average Log_10_(E/S) ratios are 0.27 ± 0.34 for PGL-3_R→K_; 0.65 ± 0.61 for PGL-3_K→R_; 0.84 ± 0.45 for PGL-3_AIDP_, and 0.98 ± 0.43 for PGL-3_AIDP+_ (**Fig. 6D**).

PGL-3_R→K_ foci are localized to exophers slightly more than WT, but not to a statistically significant difference (p=0.24). By contrast, PGL-3_K→R_ (p=0.016), PGL-3_AIDP_ (p<0.0001), and PGL-3_AIDP+_ (p<0.0001) foci localize more significantly than PGL-3_R→K_ foci to exophers (**Fig. 6D**). These data demonstrate that while PGL-3 WT is distributed nearly evenly between the soma and exopher, the PGL-3_K→R_ and PGL-3_AIDP_ variants are multi-fold enriched in the exopher compartment. PGL-3_AIDP+_ exhibits the strongest effect—nearly ten-fold enrichment—indicating that rational alterations to the PGL-3 IDR can drive selective localization into exophers.

Given the observation that some PGL-3 condensate variants, namely PGL-3_K→R_, PGL-3_AIDP_, and PGL-3_AIDP+_, are preferentially included in exophers, we next investigated which of the parameters we measured might correlate with condensate localization to the exopher compartment. We found that condensate localization to exophers correlated with slower condensate dynamics (*in vivo* D_app_, r = -0.86) (**Fig. 6E**) and higher viscoelasticity (Log of *in vitro* |G*| at 10 Hz, r = 0.88) (**Fig. 6F**). These data suggest that highly viscous and gel-like condensates are more readily recruited to exophers. Notably, recruitment did not correlate with exopher frequency or touch sensitivity, suggesting that unlike toxic Htt-polyQ128—which impairs touch sensitivity^88^, increases exopher frequency and is recruited as exopher cargo^5^—PGL-3 extrusion is not necessarily driven by its proteotoxicity (**Fig. 6G**). Furthermore, cargo selection was independent of condensate morphology such as foci size and number per cell (**Fig. 6G**).

Overall, these data support that PGL-3 variants with slower condensate dynamics and greater viscoelasticity, measured both *in vivo* and *in vitro,* are more likely to localize to large extracellular exopher vesicles. Recruitment of condensates to exophers may be influenced by a combination of factors, but among the parameters we were able to measure, we found that it was condensate dynamics (quantified by FCS *in vivo*) and material properties (quantified by VPT *in vitro*) that correlated strongly with cargo recruitment.

## Discussion

Neuronal exophers maintain cellular health through removal of protein aggregates and damaged organelles for remote degradation^5^, but the details of their cargo composition and mechanisms governing cargo recruitment remain poorly understood. Determining whether biomolecular condensates are incorporated into extracellular vesicles remains an important open question, as this could reveal previously unrecognized routes for potential spread of pathological condensates between cells. By engineering a panel of PGL-3 variants in this study, we discovered that biomolecular condensates are recruited to exophers and their material properties indeed correlate with their selection for extrusion into exophers. We found that condensates with highly viscous liquid or gel-like properties are recruited more than dynamic liquid variants, adding to the emerging understanding of how condensate properties shape biological outcomes. Through rigorously assaying the biophysics of each variant *in vitro*, we establish a mechanistic framework for interpreting the biophysics-to-phenotype relationship *in vivo*. Our *in silico* results further corroborated the mechanistic insights by resolving how distinct molecular interactions drive phase separation. We also establish *C. elegans* TRNs as an accessible model for quantitatively studying how condensate biophysics relate to neuronal proteostasis in a live animal.

Identifying the cellular pathways responsible for the outcome of why dynamically slower condensates or even aggregates are preferentially extruded into exophers remains an open question. We have previously shown that, in addition to protein aggregates, exophers contain lysosomes, aggresomes, mitochondria, and rough ER^5,8^. In many cases, these cargo organelles appear to accrue at the pre-exopher site prior to extrusion, suggesting they are actively sorted^5^. Our previous work also shows that mechanical pressure on the neuron, often from nearby eggs, can induce exopher formation^89^, supporting a mode of cargo sorting in which cellular contents not anchored to the cytoskeleton, such as diffuse cytoplasmic protein, are essentially squeezed out of the neuron. Thus, our current understanding presents a dual mode through which PGL-3 condensates could potentially enter exophers: active exopher cargo sorting and/or bulk extrusion of non-anchored cytoplasmic contents. Future studies will utilize the robust genetic toolkit in *C. elegans* to test these mechanisms through RNAi knockdown screens of motor proteins, adapter proteins, and cytoskeletal anchoring factors. If altered condensate properties affect PGL-3 association with these factors, it could explain the observed differences in recruitment. Indeed, previous studies have shown that TDP-43 RNP granules utilize kinesin1/FMRP and dynein1/Staufen1 for neuronal transport, a process that is also sensitive to the material properties of mutant TDP-43^90,91^. Furthermore, proximity labeling by TurboID has been used to identify P granule components in the *C. elegans* gonad^92^ as well as within neuronal extracellular vesicles to identify cargo components^93^. Here, TurboID could identify differential interacting partners between PGL-3 variants that may drive their cargo recruitment.

Previous studies showed autophagic clearance of condensates is dependent on condensate material properties. In yeast, liquid Ape1 condensates recruit Atg19, which promotes engulfment by isolation membrane. However, solid mutant Ape1 (P22L) interacts with Atg19 but fails to be engulfed^94^. Conversely, in *C. elegans* embryos, P granules are cleared by autophagy in somatic cells, and gel-like states induced by EPG-2 or mutations in PGL-1 increase autophagic clearance^95^. Our work shows exophers also clear condensates, such that gel-like condensates are preferentially removed over liquid condensates. Perhaps solid condensates that fail autophagic clearance are instead extruded via exophers for remote degradation.

Defining how condensate properties dictate biological outcomes in living, multicellular organisms requires developing model systems where both biophysical and phenotypic measurements can be performed. In this work, we employ FCS to quantify condensate dynamics in *C. elegans* adult neurons. FCS measurements distort with depth, and thus the transparent body of the adult worm and superficial position of TRNs—embedded directly beneath the cuticle against the coverslip— provide an ideal system for *in vivo* biophysical measurements. Our *in vivo* FCS data are supported by rigorous *in vitro* measurements of condensate dynamics via FRAP, and viscoelasticity via VPT, that show broadly consistent trends in our *in vivo* and *in vitro* data. However, the diffusivity values differ *in vivo* vs *in vitro*. This mismatch likely stems from the complex intracellular milieu and interacting partners, such as cytoplasmic mRNA, that likely influence the resulting condensate properties but are missing from the *in vitro* environment.

Our studies also highlight the importance of applying complementary techniques for side-by-side comparisons of the biophysical assays between *in vivo* experiments and minimally reconstituted *in vitro* systems. This multidisciplinary approach is emerging as a trend in the field—seen in studies of Shank3 in mouse hippocampal neurons, SPD-5 in *C. elegans* embryos, and MEC-2 in TRNs^39–41^—where condensate material properties are probed extensively *in vitro*, before being probed by the currently limited toolset of *in vivo* condensate biophysical measurements. A recurring challenge highlighted in our data and across these recent studies is the reliance on *in vivo* FRAP; because of subcellular size constraints, these systems often necessitate full-foci bleaching rather than partial bleaching. We found that *in vivo* FRAP could qualitatively capture the gel-like nature of PGL-3_AIDP+_ but lacks the resolution to quantitatively distinguish between the liquid PGL-3 variants. Our application of FCS in TRNs, and *in vivo* single-molecule tracking demonstrated in the Shank3 study^41^, present quantitative tools for measuring condensate dynamics *in vivo*. Recent advances have also made it possible to measure condensate material properties such as viscoelasticity and interfacial tension using micropipette aspiration^96,97^ and flicker spectroscopy^98^ in mammalian tissue culture and larger isolated systems such as *Xenopus* oocyte nuclei. However, to our knowledge, direct application of these techniques to small intact animal neurons such as *C. elegans* TRNs is unprecedented but may be possible in future studies.

In conclusion, we were able to apply a molecular grammar approach to engineering PGL-3 variants to achieve a wide spectrum of material properties that impact exopher cargo sorting. Furthermore, our computational simulations reveal distinct interaction patterns associated with the altered C_sat_ values and condensate material properties. Future investigation can apply this workflow to investigate disease-relevant LLPS and amyloid-forming proteins such as FUS, TDP-43, and α-Synuclein^99^, and establish exophers as a robust arm of proteostasis capable of expelling multiple neurodegenerative disease proteins beyond Htt-PolyQ shown previously^5^. Furthermore, our work holds potential applications in synthetic biology and therapeutics, including novel strategies for directing engineered proteins to exophers or loading cargo into therapeutic extracellular vesicles^100,101^. Ultimately, integration of protein engineering and live-animal imaging establishes the technical foundation to not only observe the biophysical landscape of the aging neuron but also opens avenues for future efforts to reprogram the neuronal milieu to combat disease.

## Materials and Methods

### Cloning

Transgenic *C. elegans* expression vectors were generated through Gateway Cloning. The pJW1.15 plasmid (pDONR PGL-3 ORF) was shared by Geraldine Seydoux. Barth Grant gifted a modified pCFJ1662 destination vector (Addgene #51482) containing a *mec-7* promotor, mNeonGreen with introns, in-frame Gateway site, and let858 3’UTR. This plasmid also contained hygromycin resistance for strain selection, and these sequence features were all contained within a mos1 transposable element for genomic integration. PGL-3 ORF was cloned into the modified pCFJ1662 vector using Gateway™ LR Clonase™ II Enzyme mix (Thermo #11791020) yielding pEC02 (**Table S1**).

Mutated PGL-3 IDR sequences were synthesized from Eurofins Genomics and plasmids for AIDP sequences were shared by the Kiick Lab (**Table S1**). The mutant IDR and AIDP sequences were cloned into pDONR PGL-3 replacing the wild-type IDR using Golden Gate cloning with PaqCI (NEB #R0745S). The pDONR PGL-3 variants were cloned into the modified pCFJ1662 destination vector described above yielding pEC50, pEC52, pEC61, and pEC62 (**Table S1**).

Bacterial plasmids were cloned using the backbone from Addgene Plasmid #98651, the bacterial mNeonGreen sequence from Addgene Plasmid #175947, and the PGL-3 sequences from the pDONR plasmids above. Using Golden Gate cloning, we removed MBP-FUS from p98651, while keeping the N-terminal His tag, and replaced it with mNeonGreen and PGL-3 with a TEV cleavage site in between to generate pEC56, pEC63, pEC64, pEC65, and pEC66 (**Table S1**).

The KKEE mutation (K126E, K129E) was edited into pEC02 using Golden Gate cloning to yield pEC67. Golden gate cloning was also used on pEC02 to remove the PGL-3 sequence and replace with a stop codon to yield pEC72 (**Table S1**).

### Generation of strains

ZB4065 *bzIs166 [Pmec-4::mCherry]* was used as the background for generating all other strains used in this study. ZB5916 *bzIs166[Pmec-4::mCherry]; bzSi78[Pmec-7::mNG::PGL-3 + Prps-0::HygR]* was generated by injecting pEC02 with co-injection plasmids pGH8, pCFJ90, pCFJ601, and pMA122 from Barth Grant (Jorgensen Addgene kit # 1000000031) (**Table S2**). ZB4065 worms were injected with the plasmid mix and selected for using heat-shock at 34 °C for 2 hours and 0.125mg/mL hygromycin (GoldBio #H-270-1) as previously described (Jorgensen Addgene kit # 1000000031). Similarly, pEC50, pEC52, pEC61, pEC62, pEC67, and pEC72 were injected into ZB4065 to generate mutant PGL-3 strains ZB5917, ZB5918, ZB5919, ZB5920, ZB5960, and ZB5959, respectively (**Table S2**). Once homozygosity was confirmed, animals were cultured at 20 °C and grown on standard OP50 seeded nematode growth media (NGM) without hygromycin as previously described^8^. Finally, strains were selected based on their mean fluorescence intensity within the touch neuron using fluorescence microscopy to select strains with similar expression.

### Synchronization and exopher induction

Animals were synchronized by egg lay with 20 gravid adults on OP50 seeded NGM plates for 3 hours at 20 °C as previously described^8^. Adults were removed and the plates incubated at 20 °C for 96 hours until day 2 of adulthood. In all exopher experiments, animals were transferred to unseeded NGM plates and fasted for 3 hours prior to imaging to induce exopher production^9^.

### Spinning disk confocal imaging

*In vivo* fluorescent imaging (except for FRAP time series) was captured using a Zeiss AxioObserver7 microscope with an alpha PlanApo 100x/1.46 NA oil objective equipped with X-Light V2 Spinning Disk Confocal Unit (CrestOptics), 7-line LDI Laser Launch (89 North), Photometrics Prime 95B Scientific CMOS camera, using Metamorph 7.7 software. For consistency, mCherry was imaged with the 555 nm laser line at 20% laser power with 25 ms exposure time and mNeonGreen with the 470 nm laser line at 30% laser power with 50 ms exposure time.

Animals were paralyzed in 66 mM levamisole (MPBio #02155228-CF) and mounted on 5% agarose pads with a #1.5 cover slip as previously described^8^. TRNs without an exopher event were imaged with 21 z-stack images with a step size of 0.2 μm. TRNs with an exopher event were imaged using the same step size but with as many stacks as needed to capture both the soma and exopher which are not always in the same z-plane. Images shown are maximum Z projections with brightness and contrast adjusted globally in FIJI to improve visibility of the cell and foci. Fluorescence intensity values can be found in **SI Fig. 1**.

FIJI was used to analyze images by assigning a region of interest (ROI) around the soma or exopher using the mCherry channel. Mean intensity of the ROI for each z-stack was measured, but only five brightest consecutive slices were averaged. This represents roughly the center of the object and avoids gaps that would otherwise occur at the extremes when measuring a roughly spherical object within a cylindrical ROI. The same ROI size and shape was used to measure the same five slices of the worm outside the neuron to subtract the nearby background autofluorescence intensity of the animal. When calculating Log_10_(E/S) for multiple exopher events from a single neuron, the mean intensity was calculated by taking the weighted average of the exophers considering the area of each exopher. Otherwise, Log_10_(E/S) was simply calculated using the mean intensity of the exopher and soma.

### Exopher counting

To count exophers, animals were mounted on ringed microscope slides (Avantor #48349-003) in 66 mM levamisole with 24mmx60mm #1 cover glass (Globe Sci #1419-10). Slides were imaged using an ECHO Revolve fluorescent microscope. Exophers were scored as previously described^8^.

### Touch sensitivity assay

Touch sensitivity assays were performed similarly as previously described^5,7,86^. Briefly, animals were synchronized as described above but for 72 hours until day 1 of adulthood. Day 1 adults were tested without fasting to minimize exopher formation. Animals were stroked with a gentle hair pick across the anterior portion of the body. A pause in forward crawling or complete reversal was counted as a positive response. Animals were tested 5 times each with 10 animals per strain per trial. Data in **Fig. 6C** are means of the percentages of responses out of the 50 total touches per strain per trial. Experiments were prepared blinded to avoid bias.

### Protein expression and purification

For bacterial expression, each DNA plasmid was transformed into BL21 (DE3) competent *E. coli* (New England Biolabs). The clonal culture was grown overnight at 37 °C in 25 mL sterile LB media while shaking at 250 rpm containing kanamycin (50 µg/ml). 25 mL of overnight culture media was then used to evenly inoculate 2x 500 mL of Terrific Broth medium (Fischer Scientific) supplemented with 4g/L glycerol and 50 µg/ml kanamycin for protein expression. The 500 mL cultures were grown at 37^0^C while shaking at 250 rpm until 0.6-0.8 OD and isopropyl β-D-1-thiogalactopyranoside (IPTG) was added to a final concentration of 0.5 mM to induce protein expression. After 20 hours of protein expression at 18 °C, cells were harvested by centrifugation at 4000 rpm at 20 °C for 30 min. Cell pellets were resuspended in lysis/wash buffer (1 M NaCl, 20 mM Tris, 20 mM imidazole, 1 mM DTT, EDTA-free protease inhibitor, pH 8) and lysed by sonication while on ice. Lysate was clarified by centrifugation at 24,000 x g for 40 min at 20 °C. The clarified lysate was then filtered with 0.22 µm filters.

PGL-3 WT, PGL-3_R→K_, PGL-3_K→R_, and PGL-3_AIDP_ were purified using a combination of Ni-NTA affinity and anion exchange on AKTA Pure FPLC system. In the first stage, 1 mL nickel-charged HisTrap columns (Cytiva) were used for affinity chromatography of the His-Tagged proteins. After loading the proteins onto the column, the column was washed with 15 column volumes (CV) of lysis/wash buffer (1 M NaCl, 20 mM Tris, 20 mM imidazole, 1 mM DTT, pH 8). Proteins were eluted using a 0-100% linear gradient over 30 CVs with elution buffer (1 M NaCl, 20 mM Tris, 250 mM imidazole, 1 mM DTT, pH 8). For further purification, a subset of elution fractions spanning the peak in the chromatogram were collected and dialyzed overnight using 20 kDa MWCO membranes against 500 mM NaCl, 20 mM Tris, 1 mM DTT, pH 7.5, at room temperature with at least 2 changes of buffer. FPLC flow through and remaining eluate fractions containing the desired proteins were flash frozen and preserved for future use.

For the second stage for anion-exchange chromatography, the dialyzed proteins were diluted 1:5 in 20 mM Tris, 1 mM DTT, pH 7.5 to bring the protein samples to low-salt conditions and filtered with 0.22 µm filters. The samples were then loaded onto 1 mL HiTrap Q HP column (Cytiva) and washed with 10 CVs of low salt buffer (100 mM NaCl, 20 mM Tris, 1 mM DTT, pH 7.5). Proteins were eluted using a 0-100% linear gradient over 30 CVs with high salt buffer (1 M NaCl, 20 mM Tris, 1 mM DTT, pH 7.5). Finally, the eluate spanning the chromatogram peak was transferred and dialyzed (20 kDa MWCO) into 300 mM NaCl, 20 mM Tris, 1 mM DTT, pH 7.5 for PGL-3 WT, PGL-3_R->K_, PGL-3_K->R_, and into 500 mM NaCl, 20 mM Tris,1 mM DTT, pH 7.5 for PGL-3_AIDP_.

PGL-3_AIDP+_ was purified using a 1-step Ni-NTA affinity chromatography following the above protocol and dialyzed (20 kDa MWCO) into 800 mM NaCl, 20 mM Tris, 1 mM DTT, pH 7.5. Following dialysis (anion exchange eluates for PGL-3 WT, PGL-3_R->K_, PGL-3_K->R_, PGL-3_AIDP_ and affinity eluates for PGL-3_AIDP+_), proteins were typically used directly for *in vitro* assays without snap-freezing.

The purity of samples post affinity and anion exchange chromatography elution was assessed by SDS-polyacrylamide gel electrophoresis (SDS-PAGE) using NuPAGE 4-12% Bis-Tris gels (Invitrogen) and stained using a Coomassie stain (GelCode Blue Safe Protein, Thermo Scientific) (**SI Fig. 8**).

### Microscopy for phase separation assay

Each protein sample was prepared at the desired final concentration by diluting the dialyzed stock sample at high salt with zero salt buffer (20 mM Tris, 1 mM DTT, pH 7.5) to achieve the assay buffer conditions of 150 mM NaCl, 20 mM Tris, 1 mM DTT, pH 7.5. Protein concentrations were measured based on their absorbance at 280 nm using a Nanodrop spectrophotometer (ThermoFisher). 260/280 absorbance ratio was monitored and only samples below 0.7 ratio were used for characterization.

The well plates used for microscopy were 16-well glass-bottom chambered coverslips (#1.5 glass thickness; Grace Bio-Labs) that were pretreated with 5% Pluronic F-127 (Sigma-Aldrich) for a minimum of 10 min. The coated wells were washed with assay buffer followed by transferring the protein sample to the imaging well at room temperature to initiate droplet assembly. Confocal imaging of the droplet assembly was performed at ∼1 hour on a Zeiss Axio Observer 7 inverted microscope equipped with an LSM900 laser scanning confocal module and employing a 63x/1.4 NA plan-apochromatic, oil-immersion objective. mNG was excited to fluoresce with a 488 nm laser and fluorescence images were captured using GaAsP detectors. All microscopy experiments were conducted at room temperature (19–21 °C).

### Sedimentation assay

A sedimentation assay was used to measure the saturation concentration C_sat_ of proteins by measuring the dilute phase concentration in a phase separated protein sample at 150 mM NaCl, 20 mM Tris, 1 mM DTT, pH 7.5. Samples were prepared as described for the phase separation assay and incubated for 1 hour at room temperature. The samples were then centrifuged for 20 min at 16,000xg at 20 °C in a temperature-controlled centrifuge to separate the dilute phase supernatant and dense phase pellet. The supernatant was then sampled and its absorbance at 280 nm was measured on a NanoDrop to determine protein remaining in the dilute phase to assess the C_sat_. Data presented in **Fig. 3C and 4C** are the average of two independent trials; error bars represent ± S.D.

### Video particle-tracking microrheology

Each sample was prepared at the desired final protein concentration for microrheology following the same procedure as described in the above section. Next, 120-μL protein sample was mixed with red fluorescent carboxylate-modified polystyrene beads (1 µm diameter; FluoSpheres, Invitrogen) before initiating droplet assembly in a 384-well plate (#1.5 high-performance cover glass, Cellvis). The well plate was sealed using a transparent sealing film to avoid sample evaporation. The samples were then incubated at room temperature for ∼45 mins and then observed under the microscope to verify that the tracer beads were embedded in the condensates (**Fig. 5a**). Next, the samples in the well plate were centrifuged at 300xg for 1 min to form a condensate layer or larger-sized droplets (>30 µm in diameter); the purpose of this step was to avoid boundary effects and prevent flow of the condensates. The microscope focus was adjusted to the midsection of the condensate layer for VPT acquisition.

Epifluorescence video imaging was initiated at the 1 hour timepoint using the Zeiss Axio Observer 7 inverted microscope equipped with an Axiocam 702 monochrome sCMOS camera (Zeiss), employing a x63/1.4-NA plan-apochromatic oil-immersion objective, with fluorescence excitation using a 575-nm light-emitting diode (Colibri 7; Zeiss). Videos of the tracer beads diffusing within the condensate were collected at 200 frames per second. Imaging was conducted at room temperature (19–21 °C). For each variant, at least two independent samples were made on different days, and four to five videos were collected from each sample, with each video containing ∼10–50 tracer beads. Ensemble MSDs were calculated from these tracer beads in each video. Data presented in **Fig. 5B, C** is the average of the videos from these independent trials. To employ microrheology for PGL-3_R→K_ variant, which does not phase separate at 150 mM NaCl, 20 mM Tris, 1 mM DTT, we mixed the protein sample with 4% Polyethylene Glycol (PEG) 3500 to induce phase separation. To verify if PEG 3500 significantly alters the dynamics of the condensates at the concentration used, we performed microrheology on the PGL-3 WT condensates with 4% PEG 3500. We found no significant difference in the viscosity for PGL-3 WT condensates with and without PEG (**SI Fig. 9**), showing no dramatic alteration in the dynamics of the condensates in the presence of this molecular crowder.

Data analysis to calculate the ensemble MSDs and viscosity, was conducted as described previously^48^. Next, the frequency-dependent viscoelastic moduli of the condensates were calculated from the MSD curves. The viscoelastic moduli of a material can be related to MSD of probe particles through the generalized Stokes-Einstein relation (GSER) equation:

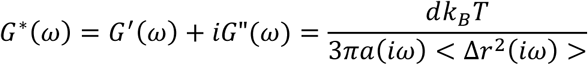

Where *G*^∗^(*ω*) is complex shear modulus, d is the number of dimensions (here d=2, because data collection and analysis were conducted in the x-y plane), k_B_ is the Boltzmann’s constant, T is the temperature, a is the bead radius, *G*^′^(*ω*) and *G*”(*ω*) are the elastic (storage) and viscous (loss) moduli, respectively and <Δ*r*^2^(*iω*)> is the Fourier transformation of the MSD <Δ*r*^2^(*t*)>. An algebraic expression proposed by Mason^85^ was used to convert the time-space MSD to frequency-space viscoelastic moduli and can be written as

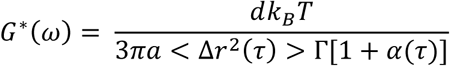

Where *α*(*τ*) = *dln*(Δ*r*^2^(*τ*))/*dln*(*τ*) is the local power index describing the logarithmic slope and Γ is the gamma function. To find local α(τ), we used a spline fit through the MSD curves to minimize errors due to local noise.

From G*(ω), we calculated G’(ω) and G” (ω) as following,

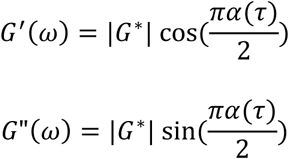

Next, spline interpolation fitting with the smoothing parameter was used to reduce the measurement noise generated in the algebraic conversion of G’ and G”. Reported viscosities and viscoelastic moduli are means ± S.D. across multiple videos (n=8-9 from at least two independent samples). Further, the loss tangent (tan δ (ω)) was calculated as a ratio of G” (ω) to G’(ω).

To remove the static error from the MSD curves for calculating viscosities and viscoelastic moduli, we corrected the ensemble-average MSD by subtracting the noise floor from the MSD curves. The noise floor of the 1 µm beads was measured by drying the beads to the glass surface of a 384-well plate. We acquired the trajectories of the beads adhered to the glass surface using the same parameters and experimental setup as used for VPT studies of the protein samples.

### Fluorescence Recovery After Photobleaching (FRAP)

FRAP experiments were performed using the same microscope and sample preparation procedure as described in the Phase Separation Assay section. A circular region of radius R = 1 μm, in the center of the protein droplets, was bleached with a 488-nm laser. Subsequent fluorescence recovery of the bleached area was recorded with a 488-nm laser for 1 min. Raw FRAP data was normalized and averaged (n = 8-9 separate droplets across two experimental setups) to obtain the final FRAP recovery curve. Normalized recovery curves were fit to an infinite boundary model to calculate the characteristic recovery half-time, τ ^75^. The apparent diffusion coefficient (D_app_) was calculated using

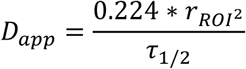

For FRAP on foci in TRNs, animals were synchronized and imaged at day 2 of adulthood and mounted on 5% agarose pads in 66 mM levamisole as described above in the spinning disk confocal section. mNG-PGL-3 variants foci in the soma were selected as ROI for bleaching and recorded with a 488-nm laser for 1 min. The fluorescence recovery from the bleached ROI was double normalized first with the whole cell fluorescent intensity to address the systematic bleaching over the experimental run, followed by normalization by setting the pre-bleach intensity to 1 and the post-bleach intensity (t=0) to 0 to compare the fluorescence recovery on a consistent scale. The normalized traces are then fit to a single exponential recovery model

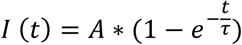

to calculate the characteristic recovery timescale, τ and percent recovery, A.

### Fluorescence Correlation Spectroscopy (FCS)

Animals were synchronized and imaged at day 2 of adulthood and mounted on 5% agarose pads in 66 mM levamisole as described above in the spinning disk confocal section. FCS was performed using a Leica TCS Stellaris8 Confocal Microscope with White Light Laser using the HC PL APO CS2 63x/1.40 NA Oil Objective. The laser was set to excitation at 488 nm and the HyD X detector set to 500 nm-550 nm emission detection with 10% gain. The pinhole was reduced to 60 μm to reduce the focal volume, and laser power was set to 0.25% for cytoplasmic measurements and reduced to 0.01-0.1% for foci measurements to prevent bleaching. Photons were collected in six 30 second segments at room temperature, taking snapshots in between to check for sample drift. Intensity fluctuations were then autocorrelated using the Leica software to get the autocorrelation function (ACF) traces of the 30 second segments. ACF traces were manually examined for the absence of spurious fluctuations at small time delays (e.g. shot noise) and absence of photobleaching. ACFs out of compliance were excluded from further analysis.

The calculated autocorrelation traces G(τ) were fitted with the model^52,55^

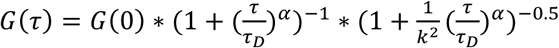

where G(0) is the value of correlation at zero lag time, τ_D_ is the translational diffusion time, α is the anomaly parameter, k is the structural parameter which is a ratio of axial to radial radii of the confocal volume, which was held fixed for the fit. Structural parameter (k) experimentally is calculated by fitting the FCS data of Alexa 488 in water (diffusivity of 435 µm^2^/s) acquired 1 µm above the coverslip. Data are fit to the autocorrelation function for simple diffusion with triplet-state kinetics to calculate the axial (r_z_) and radial radii (r_xy_) and k = r_z_/r_xy_. Values obtained for r_xy_ and r_z_ are 0.21 ± 0.01 µm and 1.72 ± 0.24 µm, respectively.

Next, the apparent diffusion coefficient (D_app_) was calculated using the equation^52,55^

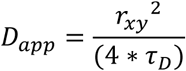

where r_xy_ is the radial radii of the confocal volume. The analysis did not focus on the fast dynamics of the fluorophore but rather on the slower translational diffusion process. Therefore, the first 1 ms of the autocorrelation data is discarded to exclude initial bleaching effect, triplet blinking effect and detector shot noise. There can be a slight difference in the actual vs calibrated values of k and r_xy_ due to refractive index mismatch and detection volume distortion using an oil immersion objective. For qualitative comparison among variants, ACF traces were normalized by setting the ACF at initial time point to 1. For data plotted in Fig. 2C, 3J, and 4K, multiple traces may originate from repeated measurements at the same focus, as well as from different foci or neurons. Therefore, n reflects the number of analyzed traces rather than the number of unique biological samples.

### Software

VPT data analysis was conducted using the open-source particle tracking package TrackPy (v0.5.0)^102^ in Python and customized as needed. Custom MATLAB code was developed to perform the algebraic conversion of the MSDs to viscoelastic moduli. Fiji (version 1.53s) was also used for image processing. JMP and Origin Pro was used for statistical testing and data plotting.

### Coarse-Grained Molecular Dynamics Simulations and Analysis

Coarse-grained (CG) phase-coexistence simulations of PGL-3 variants were performed using HOOMD-blue (versions 5.1.1 and 4.7.0)^103^, and CG single-chain simulations were carried out using LAMMPS (29 Oct 2020)^104^. Phase-coexistence simulations followed the slab-geometry protocol established in prior work^65,105–108^. Initial structures for each variant were generated with the AlphaFold server based on the corresponding sequences^109^. In all CG simulations, the two folded domains (NtDD: aa 1-210; CeDD: aa 218-447) were treated as rigid bodies with ‘fix rigid’ in LAMMPS or ‘md.constrain.Rigid’ in HOOMD-blue, while disordered regions remained fully flexible, consistent with our previously validated multidomain protein framework^107,110,111^. For contact maps, the intramolecular contacts within the folded domains were removed (set to zero), because they were constrained as rigid bodies during the simulations.

Proteins were modeled using the one-bead-per-residue HPS-Urry model^105,110^. For single-chain simulations, each construct was placed in a sufficiently large cubic box to prevent periodic self-interactions. Slab simulations containing 100 protein chains were initialized in boxes measuring 250 × 250 × 1750 Å^3^ for full-length proteins and 170 × 170 × 1190 Å^3^ for IDR-only constructs.

All CG simulations were run in the canonical ensemble (NVT) at 300 K using a 10-fs time step. Temperature was controlled using a Langevin thermostat, with a damping factor τ of 1000 ps for slab simulations and 1000 fs for single-chain simulations. Electrostatic screening was modeled implicitly using a Debye-Hückel potential with a Debye length of 1 nm, corresponding to an ionic strength of ∼100 mM. Radius of gyration, intrachain distance, and contact analyses were performed following the protocol described in our previous studies^107,108,110^.

## Acknowledgements

We thank Barth Grant, Geraline Seydoux, and Kristi Kiick for providing plasmids used for cloning. The graphical abstract (https://BioRender.com/zbzxoyj), Fig. 1A (https://BioRender.com/x0li0cs), Fig. 1F (https://BioRender.com/bj1ghl3), and Fig. 2B (https://BioRender.com/iuj0edj) were created in BioRender. Our work was supported by NIH grants 3K12GM093854 (EC), 5T32EB005583 (EC), R37AG56510 (MD), R01AG084206 (MD), R35GM142903 (BSS), and R35GM153388 (JM). We gratefully acknowledge the computational resources provided by the Texas A&M High Performance Research Computing (HPRC).

## Author Contributions

EC, MB, MD and BSS conceived the research project and designed the sequences. EC and MB performed molecular cloning to design protein sequences. MB expressed, purified all proteins, performed *in vitro* assays including sedimentation, microscopy, FRAP and microrheology. EC performed molecular cloning and microinjection to generate transgenic strains and performed *in vivo* assays including microscopy, exophers counting, touch assay, and FRAP. EC and MB performed *in vivo* FCS assay. MB analyzed data related to *in vitro* assays, *in vivo* FRAP and FCS. JM and SLW conceived CG-MD simulations. SLW performed investigation and formal analysis of CG-MD simulations. MB, EC, MD and BSS wrote the original draft. All authors reviewed and edited the manuscript.

## Generative AI declaration

Gemini was used to digitize a hand-drawn diagram of a neuron and exopher which was uploaded to BioRender and used as an asset within the graphical abstract. The original text of this manuscript was written by the authors. Gemini and NotebookLM were used to give suggestions to improve grammar, clarity, and brevity of individual sentences. All AI outputs were carefully reviewed and further edited by the authors before any changes were applied to the text to ensure the original meaning and intent were not altered by AI.

