## Supplemental Information for "Condensate Material Properties Influence Cargo Selection for Neuronal Extrusion via Large Extracellular Exopher Vesicles"

**Supplementary Information**

**Supplementary Figures**

**SI Fig. 1**

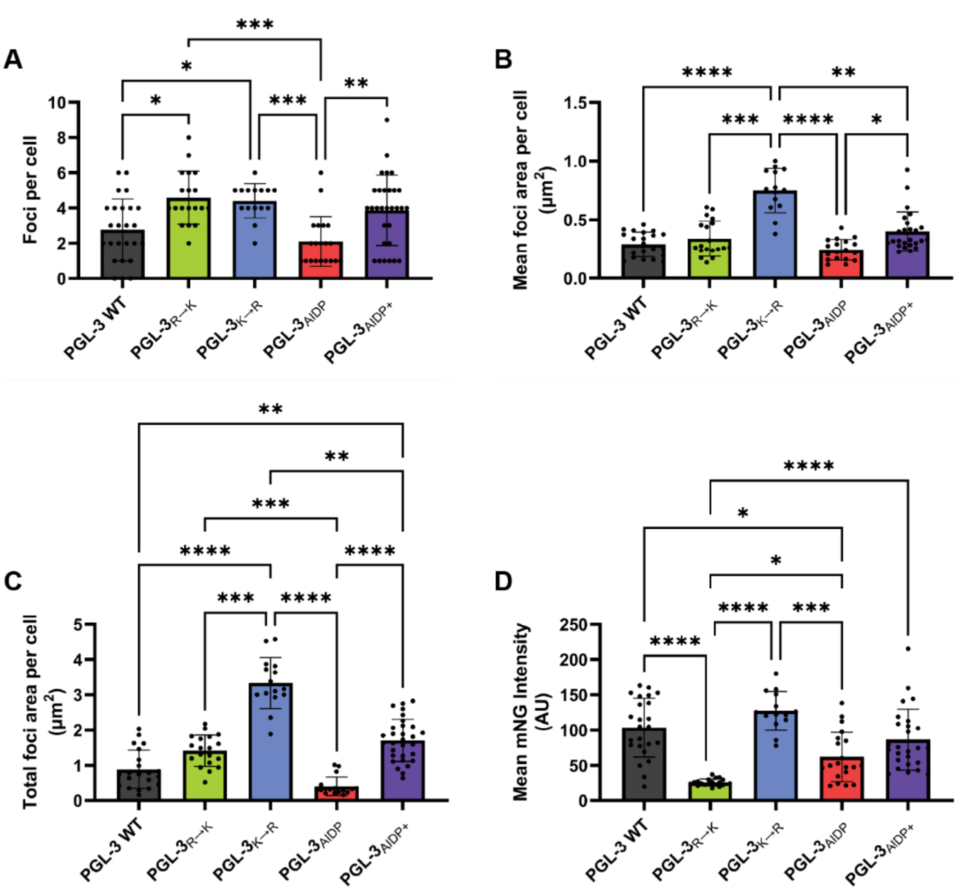


**SI Fig. 1:** Quantification of foci properties in touch neurons expressing mNG-PGL-3 variants. **(A)** Number of foci per cell soma, **(B)** average area of all foci in each cell soma, **(C)** total area of all foci in each cell soma, **(D)** mean mNG intensity across the entire cell soma. All animals were adult day two, neurons imaged were a mix of ALMR and ALML. N = 14-26 neurons for each variant. Statistics are Kruskal-Wallis test with Dunn's multiple comparisons test (* $p<0.05$, ** $p<0.01$, *** $p<0.001$, **** $p<0.0001$).

**SI Fig. 2**

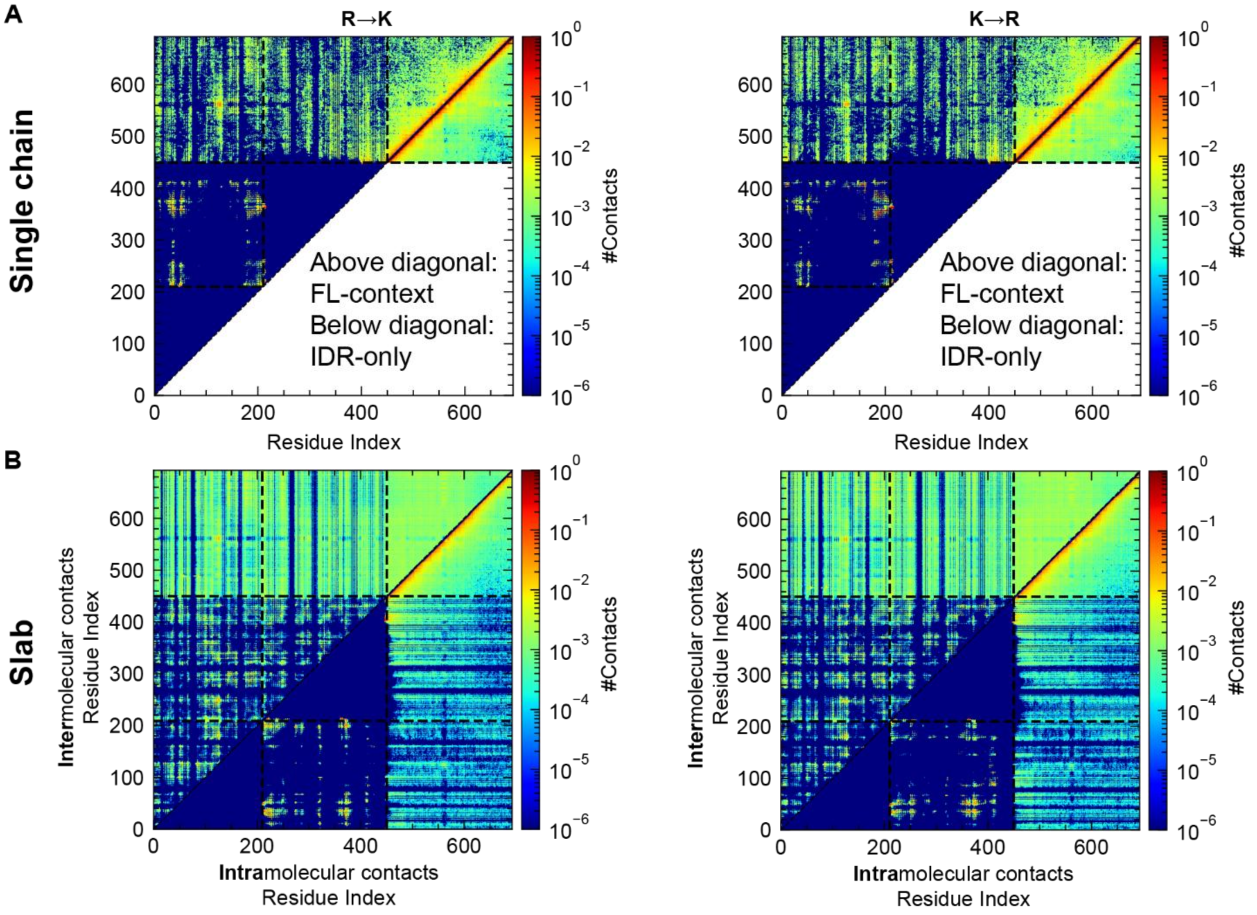


**SI Fig. 2: (A)** Intramolecular contact maps from single-chain simulations of full-length (FL, above diagonal) and IDR-only (below diagonal) constructs, representing the dilute-phase ensemble. Dashed lines delineate domain boundaries domain (NtDD, AA 1-210; CeDD, AA 218-450; IDR, AA 451-693). Contacts within folded domains are omitted (set to zero) due to rigid-body constraints applied during the simulation. **(B)** Condensed-phase contact maps derived from slab simulations. The upper-left and lower-right triangles of each matrix display intermolecular (chain–chain) and intramolecular (within-chain) contacts, respectively. Domain boundaries and rigid-body constraints match those in panel **A**.

**SI Fig. 3**

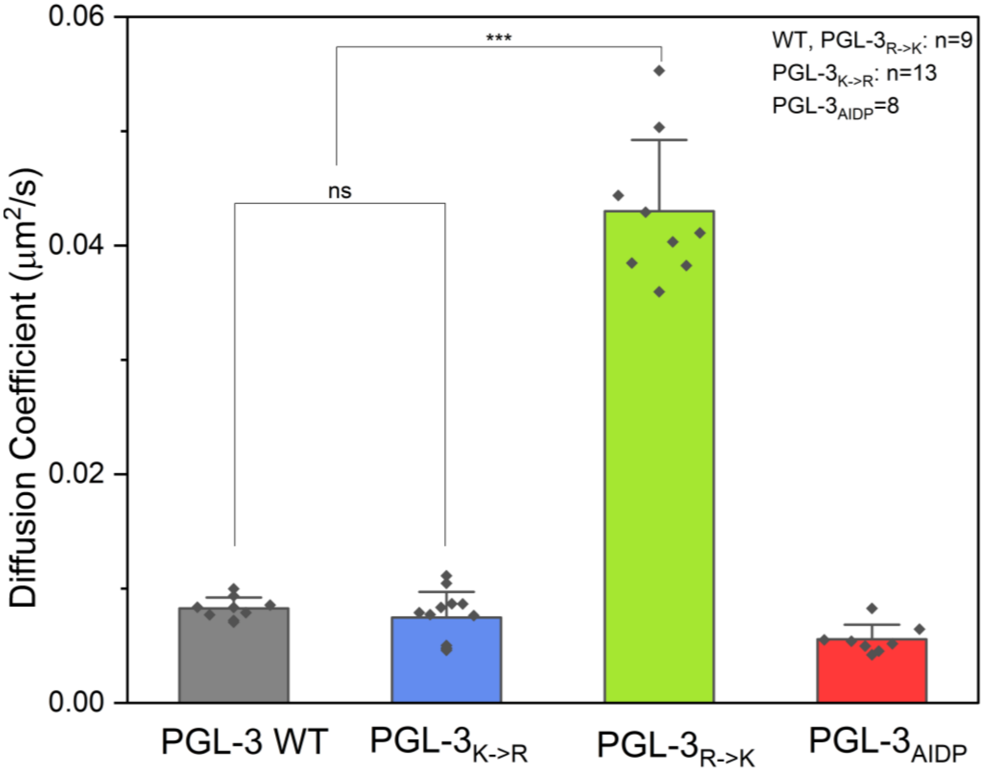


**SI Fig. 3:** Apparent diffusion coefficients, $D_{app}$, calculated by fitting the FRAP recovery traces from **Fig. 3G and Fig. 4H** to an infinite boundary model in two dimensions (n=8-9 from two independent experiments for PGL-3 WT, PGL-$3_{R\rightarrow K}$ and PGL-$3_{AIDP}$, n=13 for PGL-$3_{K\rightarrow R}$ from 3 independent experiments).

**SI Fig. 4**

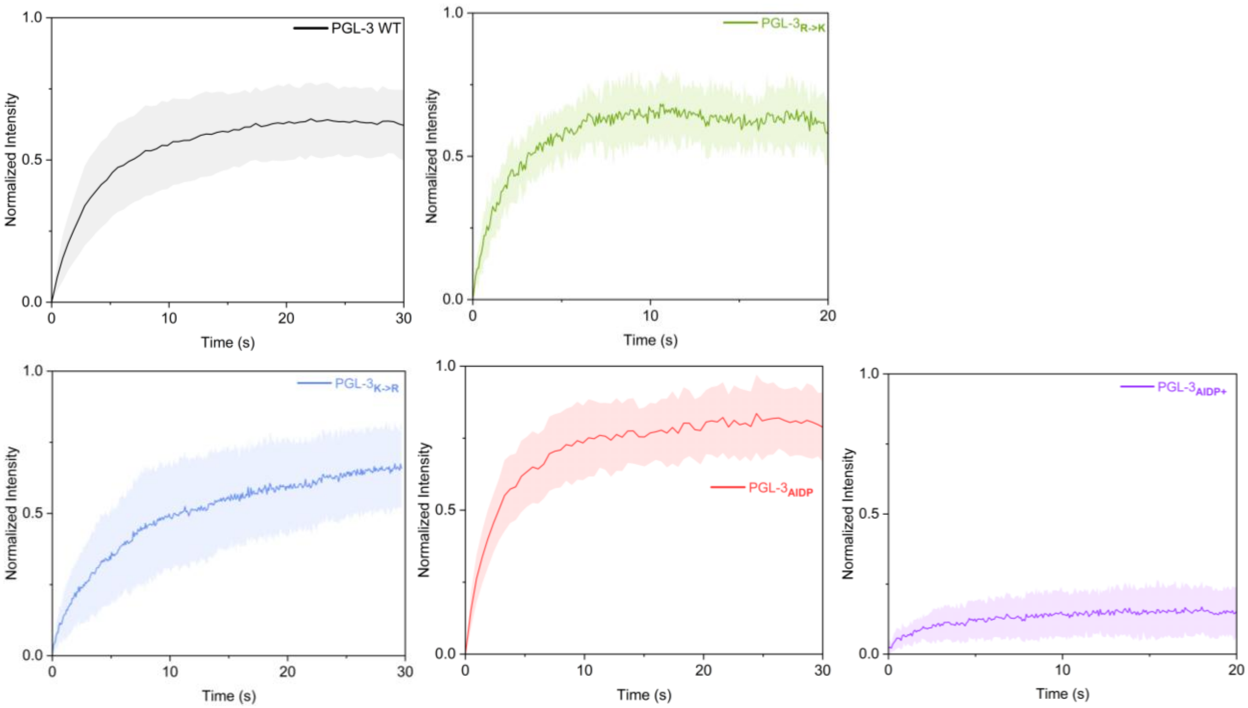


**SI Fig. 4:** *In vivo* whole foci FRAP traces for data in **Fig. 3I and 4J**. Y-axis represents normalized intensity where 0 is the bleached ROI intensity and 1 is the pre-bleached ROI intensity. Data are presented as mean $\pm$ S.D. (PGL-3 WT: *n*=19, PGL-3$_{R\rightarrow K}$: *n*=8, PGL-3$_{K\rightarrow R}$: *n*=7, PGL-3$_{AIDP}$: *n*=12, PGL-3$_{AIDP+}$: *n*=10).

**SI Fig. 5**

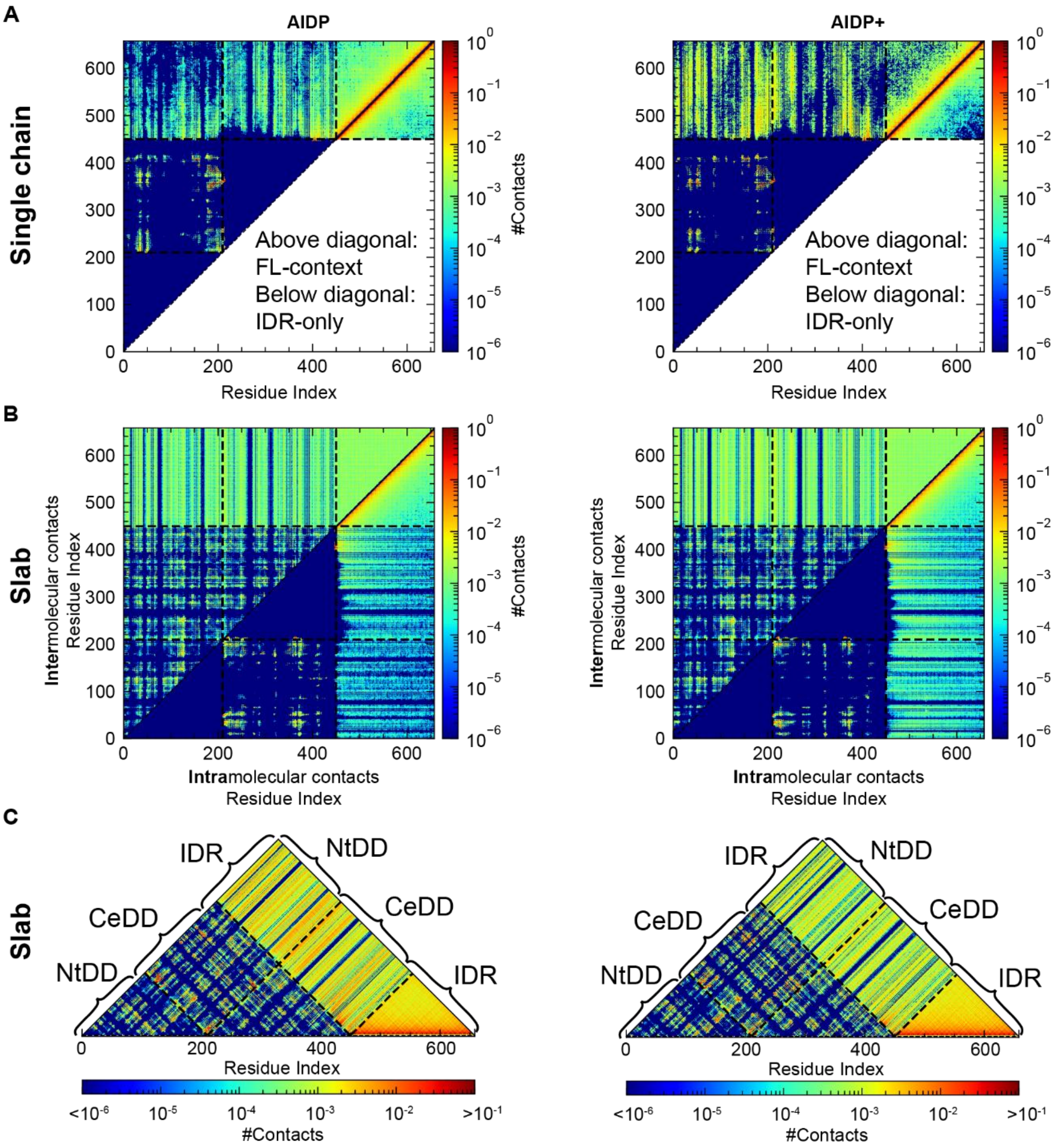


**SI Fig. 5: (A)** Intramolecular contact maps from single-chain simulations of full-length (FL, above diagonal) and IDR-only (below diagonal) constructs, representing the dilute-phase ensemble. Dashed lines delineate domain boundaries. Contacts within folded domains are omitted (set to zero) due to rigid-body constraints applied during the simulation. **(B)** Condensed-phase contact maps derived from slab simulations. The upper-left and lower-right triangles of each matrix display intermolecular (chain–chain) and intramolecular (within-chain) contacts, respectively. Domain boundaries and rigid-body constraints match those in panel **A**. **(C)** Combined contact map integrating inter- and intramolecular interactions within the condensed phase.

**SI Fig. 6**

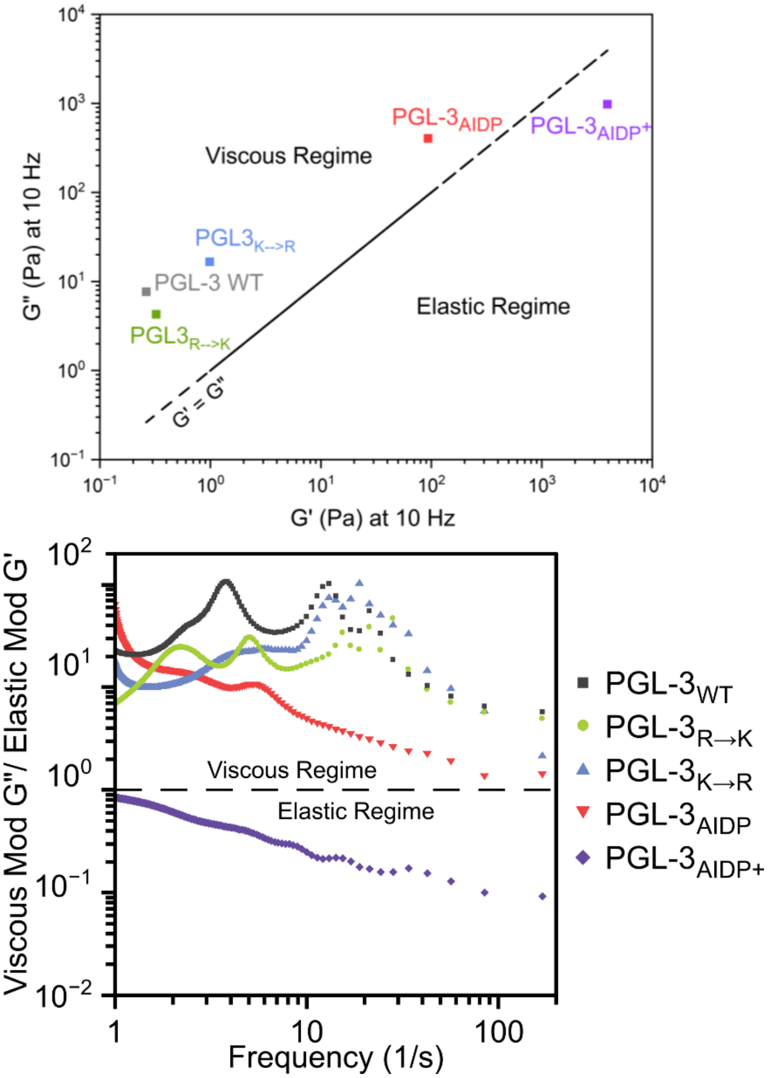


**SI Fig. 6 (A)** A material state diagram based on measured viscoelastic moduli at 10Hz. A reference line at G' = G" to delineate dominantly viscous regime versus dominantly elastic regime. **(B)** Loss tangent, defined as the ratio of viscous modulus G" to elastic modulus G', plotted against the experimental probed frequency for condensates formed by all PGL-3 variants. Dotted line represents the delineation between dominantly viscous and dominantly elastic regime which confirmed PGL-$3_{AIDP+}$ condensates show dominantly elastic behavior in the experimental probed frequencies. (n=8-9 from two independent experiments for all variants). We also observed that variants show an increase in condensate viscoelasticity in the order PGL-$3_{AIDP+}$ > PGL-$3_{AIDP}$ > PGL-$3_{K\rightarrow R}$ > PGL-3 WT ≈ PGL-$3_{R\rightarrow K}$, suggesting an important role of interdomain interactions in strengthening the intra-condensate network, thereby impacting viscoelasticity.

**SI Fig. 7**

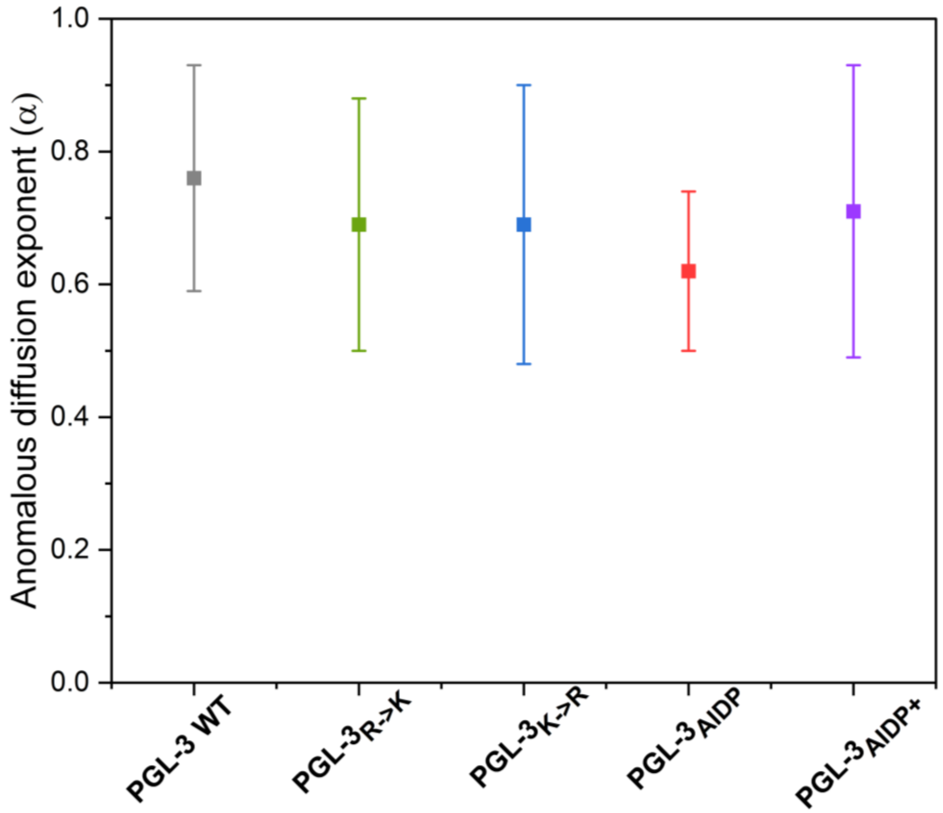


**SI Fig. 7:** Anomalous diffusion exponent (α) obtained from the model fit of ACFs from **Fig. 3J & Fig. 4K** confirming diffusion of protein molecules of the PGL-3 variants in the foci is anomalous.

**SI Fig. 8**

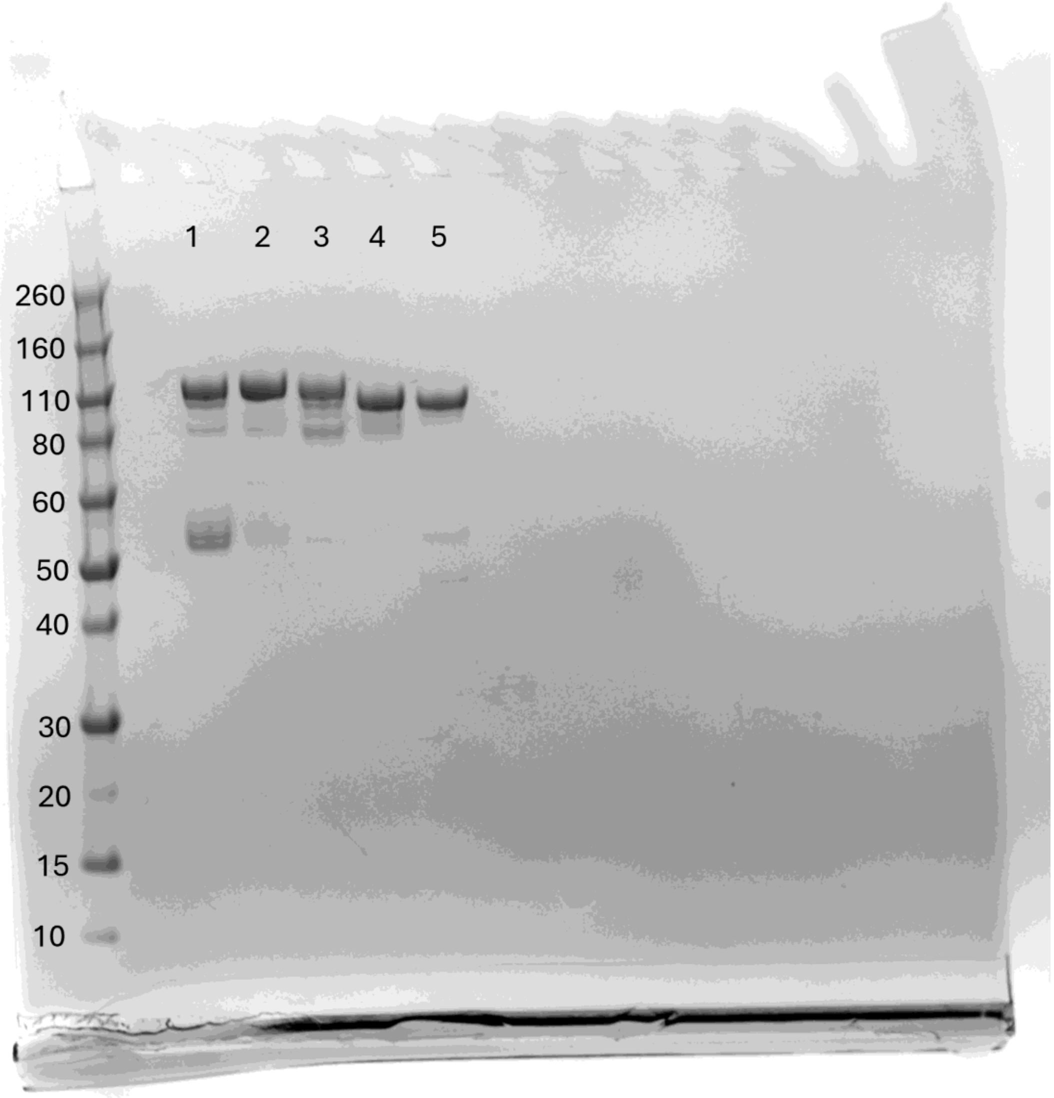


1. PGL-3 WT 2. PGL-3$_{R \rightarrow K}$ 3. PGL-3$_{K \rightarrow R}$4. PGL-3$_{AIDP}$ 5. PGL-3$_{AIDP+}$

**SI Fig. 8.** SDS-PAGE of all sequences showing a unique band after purification. All samples were loaded at 2 µg and run under standard protocols. Representative image displays results from all samples in a single run. However, SDS-PAGE gel on individual samples were run separately during purification steps. Slight variations in the MW of the constructs arise not only from variations in their theoretical MW but also likely variations in their charge/mass ratios because of their differential adsorption of SDS due to differences in their ionic character.

**SI Fig. 9**

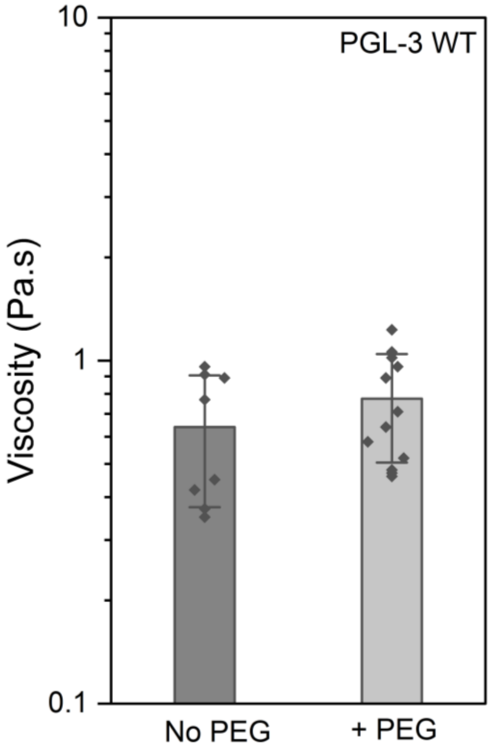


**SI Fig. 9:** Viscosity calculated from VPT for PGL-3 WT with and without 4% PEG 3500 showing no significant effect of molecular crowder on the viscosity of the condensates.

**Table S1: Plasmids**

| Plasmid | Gene of interest | Backbone from |
|---|---|---|
| pEC02 | *Pmec-7::mNG::PGL-3* | PCFJ1662 (Plasmid #51482) |
| pEC50 | *Pmec-7::mNG::PGL-3 (R485K, R579K, R627K, R634K, R638K, R641K, R648K, R650K, R658K, R661K, R665K, R668K, R676K, R682K, R690K)* | PCFJ1662 (Plasmid #51482) |
| pEC52 | *Pmec-7::mNG::PGL-3 (K458R, K480R, K503R, K520R, K525R, K548R, K555R, K571R, K602R)* | PCFJ1662 (Plasmid #51482) |
| pEC61 | *Pmec-7::mNG::PGL-3$_{1\text{-}450}$:: RSEFNG::(GRGDSPYS)$_{25}$* | PCFJ1662 (Plasmid #51482) |
| pEC62 | *Pmec-7::mNG::PGL-3$_{1\text{-}450}$:: RSEFDP::(GRGNSPYS)$_{25}$* | PCFJ1662 (Plasmid #51482) |
| pEC67 | *Pmec-7::mNG::PGL-3 (K126E, K129E)* | PCFJ1662 (Plasmid #51482) |
| pEC72 | *Pmec-7::mNG* | PCFJ1662 (Plasmid #51482) |
| pEC56 | *His6::mNG::TEVsite::PGL-3* | MBP-FUS_FL_WT (Plasmid #98651) |
| pEC63 | *His6::mNG::TEVsite::PGL-3 (R485K, R579K, R627K, R634K, R638K, R641K, R648K, R650K, R658K, R661K, R665K, R668K, R676K, R682K, R690K)* | MBP-FUS_FL_WT (Plasmid #98651) |
| pEC64 | *His6::mNG::TEVsite::PGL-3 (K458R, K480R, K503R, K520R, K525R, K548R, K555R, K571R, K602R)* | MBP-FUS_FL_WT (Plasmid #98651) |
| pEC65 | *His6::mNG::TEVsite::PGL-3$_{1\text{-}450}$:: RSEFNG::(GRGDSPYS)$_{25}$* | MBP-FUS_FL_WT (Plasmid #98651) |
| pEC66 | *His6::mNG::TEVsite::PGL-3$_{1\text{-}450}$:: RSEFDP::(GRGNSPYS)$_{25}$* | MBP-FUS_FL_WT (Plasmid #98651) |

**Table S2: Strains**

| Strain | Genotype |
|---|---|
| ZB4065 | *bzIs166[Pmec-4::mCherry]* |
| ZB5916 | *bzIs166[Pmec-4::mCherry]; bzSi78[Pmec-7::mNG::PGL-3]* |
| ZB5917 | *bzIs166[Pmec-4::mCherry]; bzSi79[Pmec-7::mNG::PGL-3 (R485K, R579K, R627K, R634K, R638K, R641K, R648K, R650K, R658K, R661K, R665K, R668K, R676K, R682K, R690K)]* |
| ZB5918 | *bzIs166[Pmec-4::mCherry]; bzSi80[Pmec-7::mNG::PGL-3 (K458R, K480R, K503R, K520R, K525R, K548R, K555R, K571R, K602R)]* |
| ZB5919 | *bzIs166[Pmec-4::mCherry]; bzSi81[Pmec-7::mNG::PGL-3$_{1\text{-}450}$:: RSEFNG::(GRGDSPYS)$_{25}$]* |
| ZB5920 | *bzIs166[Pmec-4::mCherry]; bzSi82[Pmec-7::mNG::PGL-3$_{1\text{-}450}$:: RSEFDP::(GRGNSPYS)$_{25}$]* |
| ZB5959 | *bzIs166[Pmec-4::mCherry]; bzSi83[Pmec-7::mNG]* |
| ZB5960 | *bzIs166[Pmec-4::mCherry]; bzSi84[Pmec-7::mNG::PGL-3 (K126E, K129E)]* |
